# Diel remodeling and cellular integration of the nitroplast

**DOI:** 10.64898/2026.04.11.717942

**Authors:** Ethan H Li, Tyler Coale, Gaëlle Toullec, Artur Czajkowski, Pierre Henri Jouneau, Tim M. Dederichs, Chandni Bhickta, Zachary Raymond, Kyoko Hagino, Vitor Hugo Balasco Serrão, Jotham Austin, Wanda M. Figueroa-Cuilan, Yannick Schwab, Francisco M. Cornejo-Castillo, Jonathan Zehr, Johan Decelle, Mohammed Kaplan

## Abstract

Nitrogen-fixing eukaryotes were not believed to exist in nature until the recent discovery of a N_2_-fixing organelle, or nitroplast, in the marine microalga *Braarudosphaera bigelowii*. This nitroplast (formerly known as UCYN-A2) has long been recognized as key cyanobacterial contributor to global oceanic N₂ fixation. However, how this novel organelle is integrated and regulated within the architecture of a eukaryotic cell remains unclear. Here, we combine multiscale volumetric imaging with cryo–electron tomography to resolve the native architecture, cellular integration, and diel remodeling of the nitroplast in cultured and environmental cells. We find that the nitroplast occupies up to 10% of the cell volume and exhibits close interfaces with multiple host organelles through membrane contact sites, while integration of this metabolically demanding compartment does not disrupt global scaling of host organelles. Interestingly, the chloroplast-to-nitroplast volume ratio is conserved across distinct life stages. Cryo-electron tomography reveals that the nitroplast retains a reinforced four-layer cyanobacterial envelope and is additionally surrounded by two host-derived layers that remodel across the day–night cycle. During daytime N₂ fixation, these host-derived barriers become locally discontinuous and the organelle interface becomes enriched with two distinct vesicle populations. Our findings suggest that dynamic control of organelle accessibility through transient membrane gating represents a fundamental strategy by which eukaryotic cells could domesticate new endosymbiotic functions during early organellogenesis.

## Introduction

Marine phytoplankton, including microalgae and cyanobacteria, are responsible for a large fraction of global photosynthesis on Earth, and therefore play a critical role in the global carbon cycle^1^. Some genera of cyanobacteria also play a role in the global nitrogen (N) cycle, as N_2_-fixers (diazotrophs) that reduce atmospheric N_2_ to biologically available ammonia^2^. While photosynthetic carbon fixation^3^ (chloroplasts) and aerobic respiration^4^ (mitochondria) were acquired early in the evolution of eukaryotes through multiple endosymbiotic events^5–9^, N_2_ fixation has long been considered restricted to prokaryotes, or eukaryotes via symbiosis with diazotrophic prokaryotes^10–12^. This raises a central question in cell evolution: can a eukaryotic cell domesticate a nitrogen-fixing bacterium into a fully integrated organelle, and if so, how is such a metabolically demanding and oxygen-sensitive machine structurally accommodated and regulated within the eukaryotic cell? These questions have therefore remained unresolved in cell and evolutionary biology.

The recent discovery of the N_2_-fixing organelle, the nitroplast^13,14^, in the marine microalga *Braarudosphaera bigelowii* (haptophyte) fundamentally changed this view. The nitroplast (formerly known as *Candidatus* Atelocyanobacterium thalassa or UCYN-A), and its close relatives have long been recognized as globally distributed cyanobacteria^15^ that contribute substantially to oceanic N₂ fixation, supplying ∼20% of fixed N_2_ in the tropical North Atlantic^16,17^. The nitroplast exhibits hallmark features of organelles, including a highly reduced genome, metabolic dependency on the host^18,19^, nuclear-encoded protein import, and synchronized division with the host cell^13^. Compared to many other known organelles, the nitroplast represents a recent evolutionary innovation, estimated to have evolved ∼100 million years ago^20,21^. As such, it provides a unique opportunity to investigate how a newly internalized metabolic compartment becomes structurally integrated and regulated within a eukaryotic cell that already contains other endosymbiotic-derived organelles such as mitochondria and chloroplasts.

Another hallmark of nitroplast integration is the coordination of its gene expression with the host cell. Nitroplast gene expression follows diel oscillations^22,23^ likely in synchrony with the host cellular rhythms, similar to other organelles whose function is coupled to the circadian cycle^24–26^. Consistent with this temporal regulation, isotope tracer experiments have shown that N_2_ fixation occurs exclusively during the daytime^27^. Accordingly, the absence of light suppresses N_2_ fixation, suggesting that the process relies on carbon derived from host photosynthesis^28^. This coupling between oxygen-sensitive N_2_ fixation and photosynthetic metabolism has also been observed in other N_2_-fixing endosymbionts^29^.

Despite its ecological significance in the global N cycle of the ocean, our current understanding of nitroplast structure remains limited. Previous studies relied primarily on conventional microscopy or X-ray tomography^13,14,30–33^. While informative, these approaches require chemical fixation, dehydration, and staining steps that can disrupt macromolecular and membrane organization. These studies described the nitroplast as a spherical organelle (∼2 µm diameter) containing multiple thylakoid-like membranes and suggested an envelope composed of three layers corresponding to inner membrane (IM), peptidoglycan (PG), and outer membrane (OM)^30^. The organelle was further suggested to be enclosed by a host-derived membrane (HDM)^30,31^. However, the native organization of this boundary, how the nitroplast interfaces with host organelles, how metabolites are exchanged, and how diel regulation is structurally accommodated all remain unresolved.

Here, we combine multiscale volumetric imaging with *in situ* cryo–electron tomography (cryo-ET)^34^ to examine how the nitroplast is structurally integrated within its host cell and dynamically regulated across the day–night cycle. We resolve the cellular architecture of *B. bigelowii* in 3D across different life stages to assess how acquisition of the nitroplast affects scaling and spatial organization of host organelles, and determine the native macromolecular organization of the nitroplast during distinct diel phases revealing pronounced structural remodeling associated with daytime N_2_ fixation. Together, these analyses provide a multiscale view of how a newly acquired, energy-intensive and oxygen-sensitive compartment is accommodated within a photosynthetic eukaryotic cell, offering direct insight into the structural principles underlying early organellogenesis.

## Results

### Nitroplast integration preserves host organelle scaling and cellular architecture

To determine how acquisition of the nitroplast impacts host cell organization, we used focused ion beam–scanning electron microscopy (FIB–SEM)^35^ to generate three-dimensional (3D) reconstructions of entire cryo-fixed, freeze-substituted *B. bigelowii* cells at 8 nm resolution. The recency of nitroplast evolution provides a unique opportunity to assess whether integration of a new metabolic compartment perturbs the size, scaling, or spatial organization of pre-existing organelles in a host cell.

We reconstructed three motile *B. bigelowii* cells maintained in culture (average volume 130.5 ± 8.2 µm^3^; Movies S1-S3) and segmented the main organelles and compartments, including chloroplasts, mitochondria, nucleus, vacuoles, and the nitroplast (Fig. 1). The two plastids were the largest organelles, occupying 35% of total cell volume (Fig. S1; Table S1). The nitroplast and an electron-dense vacuole were the second largest organelles, occupying 10.2 ± 0.5% and 11.5 ± 2.2% of the cell volume, respectively, followed by the nucleus (6.5 ± 0.2%), and mitochondria (4 ± 0.2%, Fig. S1; Table S1). 3D reconstructions also revealed extensive proximity between the nitroplast and mitochondria, plastids, the nucleus, and a yet-unidentified membrane-bound compartment consisting of a branched tubular network (Fig. S2, Movie S3). These repeated physical juxtapositions in different cells suggest coordinated metabolic interactions rather than random spatial associations.

**Figure 1.**
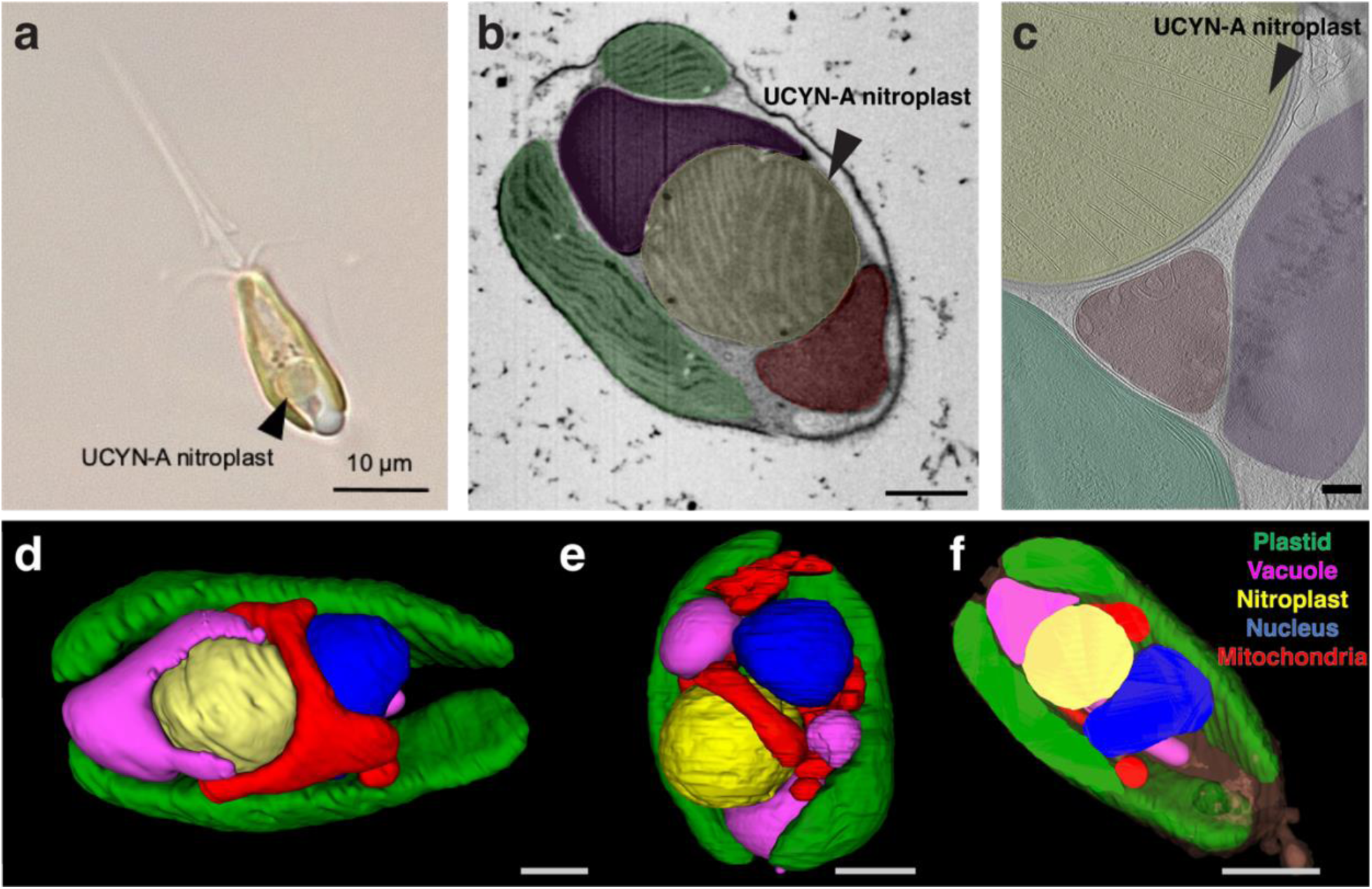
Cellular architecture of the haptophyte *B. bigelowii* harboring the nitroplast revealed by 3D electron microscopy. a, A motile *B. bigelowii* cell and nitroplast as visualized with light microscopy. **b,** A representative slice from the FIB–SEM volume illustrating organelle ultrastructure and the close spatial proximity of the nitroplast to plastids and mitochondria. **c,** A Slice through a cryo-electron tomogram of a cryo-FIB–milled *B. bigelowii* cell showing the nitroplast in a vitrified, hydrated state and illustrating the multiscale imaging workflow used in this study (light microscopy, FIB–SEM, cryo-ET). Scale bar: 10 µm (a), 2 µm (b), 200 nm (c). **d-f,** FIB–SEM–based 3D reconstructions of two cultured, motile *B. bigelowii* cells showing plastids (green), mitochondria (red), vacuole (purple), nucleus (blue), and nitroplast (yellow). Scale bar: 2 µm.

Comparison with two closely related motile haptophytes in culture lacking a nitroplast, (*Gephyrocapsa huxleyi*^36,37^ and *Phaeocystis cordata*^38^) revealed remarkably conserved proportional volume occupancy of plastids (∼31–35 %) and mitochondria (∼4–7.5 %) despite interspecies cell size differences. Thus, in these motile *B. bigelowii* cells, incorporation of the nitroplast does not measurably alter the gross proportional investment in host energy-producing organelles, suggesting that N_2_ fixation is accommodated without large-scale restructuring of cellular bioenergetics. Of note, the relative volume of the nucleus was modestly reduced in *B. bigelowii* (6.5%) compared to the other two haptophytes (∼9.3%).

We also had the opportunity to investigate the cellular architecture of *B. bigelowii* cells and their nitroplast in a different life stage, a calcified non-motile form obtained directly from the marine environment (Materials and Methods). This observation is a challenge since this form is not in culture and is considered to be possibly a resting stage, although such states are not known from other coccolithophores^39^. We therefore implemented a specific imaging workflow^40^ in order to target specifically the calcified non-motile cells with FIB-SEM from a complex environmental plankton community (Materials and Methods).

We successfully imaged the calcified form of three *B. bigelowii* cells from their natural environment (Fig. S3 and Movie S4). Compared to the non-calcified form, these cells were surrounded by thick penthaliths and exhibited approximately fourfold larger cell volumes (570.2 ± 84.6 µm^3^; Table S1). They display four thin, plate-like plastids (each ranging from 14 to 24 µm^3^) positioned at the cell periphery, consistent with previous observations^30^. The relative plastid volume occupancy was lower than in the uncalcified form (13% vs 35%). In these cells, we also observed two nitroplasts (each 7-16 µm^3^ in volume), together occupying 3.9 ± 0.7% of the cell volume, compared to ∼10% for the single nitroplast in uncalcified cells. Notably, despite substantial differences in absolute cell size and morphology between calcified and non-calcified life stages, the ratio of chloroplast to nitroplast volume remained conserved (∼3.2 in both forms).

### Cryo-ET reveals native nanoscale architecture of the nitroplast

To visualize the nitroplast in a native, hydrated state, we performed cryo-ET on plunge-frozen *B. bigelowii* cells. Because these cells (5–10 µm thick) exceed the penetration limit of electrons, we used cryo–focused ion beam (cryo-FIB) milling to generate ∼200 nm lamellae suitable for cryo-ET (Fig. S4). N_2_ fixation in *B. bigelowii* is light dependent, initiating shortly after dawn. We therefore focused first on cells collected after 1 h of light exposure (“morning nitroplasts”). We acquired 28 tomograms from 14 independent nitroplasts (Movies S5–S6). *In situ*, nitroplasts appeared spherical or ellipsoidal (2–4 µm diameter) and contained numerous internal thylakoid-like membranes extending into the matrix, also visible with FIB-SEM (Figs. 1&2a–c). These internal membranes, derived from cyanobacterial thylakoids but lacking photosystem II proteins^41,42^, varied widely in thickness (∼10–160 nm), with two dominant populations centered around ∼20 and ∼35 nm (Fig. S5).

Within the nitroplast matrix, we identified ∼65 nm globular structures attached to the external surface of thylakoid membranes (Fig. 2d,e). Their morphology and membrane association closely resemble plastoglobules in plastids^43^, suggesting conserved lipid or cofactor storage functions. Additionally, some thylakoid membranes exhibited arrays of peripheral densities reminiscent of ATP synthase assemblies on mitochondrial cristae^44^ (Fig. 2f), potentially reflecting the high energetic demand associated with nitrogenase activity. We also observed amorphous electron-dense bodies (50–150 nm) enriched near the inner membrane of the nitroplast (Fig. 2g; Fig. S6), as well as ribosomes and occasional internal vesicles (Fig. 2a,b, yellow and light blue arrows). Together, these features indicate a metabolically active matrix with complex internal organization. Another feature of morning nitroplasts was a triangular (hat-like), membrane-anchored structure protruding into the lumen of thylakoid membranes (Fig. 2h,i). We identified 25 such complexes across datasets, suggesting they are an abundant structure in the morning nitroplast.

**Figure 2.**
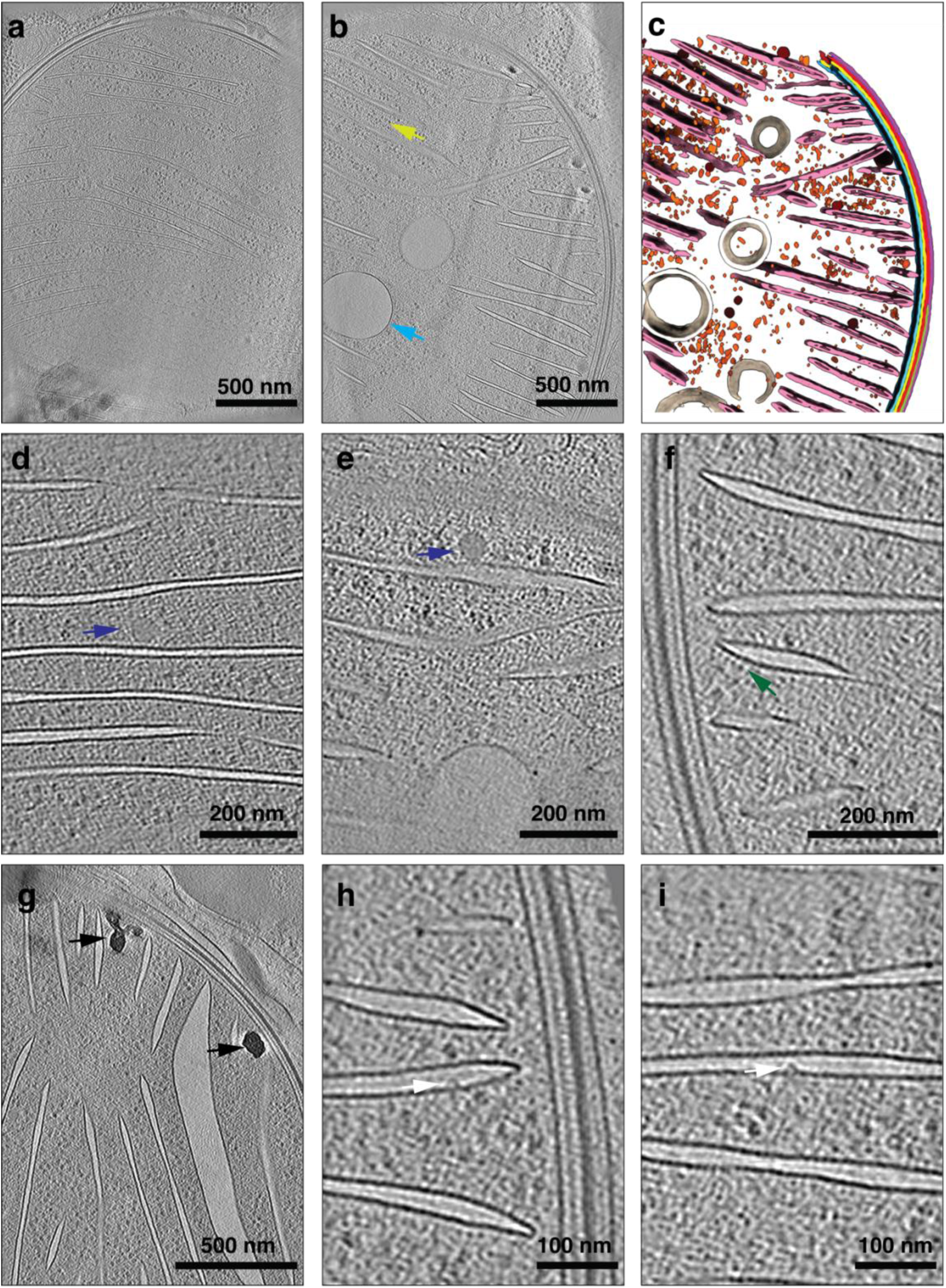
Nanoscale architecture of the morning nitroplast revealed by cryo-ET. a,b, Representative slices through cryo-electron tomograms of morning nitroplasts. **c,** 3D segmentation of the tomogram shown in **b**. **d–i,** Tomographic slices highlighting key ultrastructural features of the morning nitroplast, including plastoglobule-like structures associated with thylakoid membranes (**d,e;** blue arrows), arrays of peripheral densities on the cytosolic face of thylakoid membranes (**f;** green arrow), amorphous electron-dense bodies near the inner membrane (**g;** black arrows), and a triangular thylakoid-associated complex protruding into the thylakoid lumen (**h,i;** white arrows).

### The nitroplast is enclosed by a composite, six-layer envelope

The resolution of our cryo-tomograms revealed unexpected complexity in the nitroplast envelope. Contrary to previous interpretations from conventional transmission electron microscopy on resin-embedded cells which assigned three layers to the nitroplast envelope^30^, our results revealed that the nitroplast envelope comprises four distinct layers (Fig. 3a,b; Movie S7). Three correspond to the canonical cyanobacterial IM, PG layer, and OM. In addition, we identified a granular fourth layer (GL) of unknown composition positioned between the PG layer and the OM. Interestingly, the OM was conspicuously thick (∼12 nm), suggesting a reinforced barrier (Fig. 3). Beyond this envelope, the nitroplast was surrounded by two additional host-derived layers: a host-derived membrane (HDM) and an underlying granular layer that remained closely anchored to it (Fig. 3a,b, henceforth referred to as host granular layer “HGL”). Where the HDM drifted locally, the HGL followed, indicating that it is physically anchored to the HDM (Fig. S7a,b).

**Figure 3.**
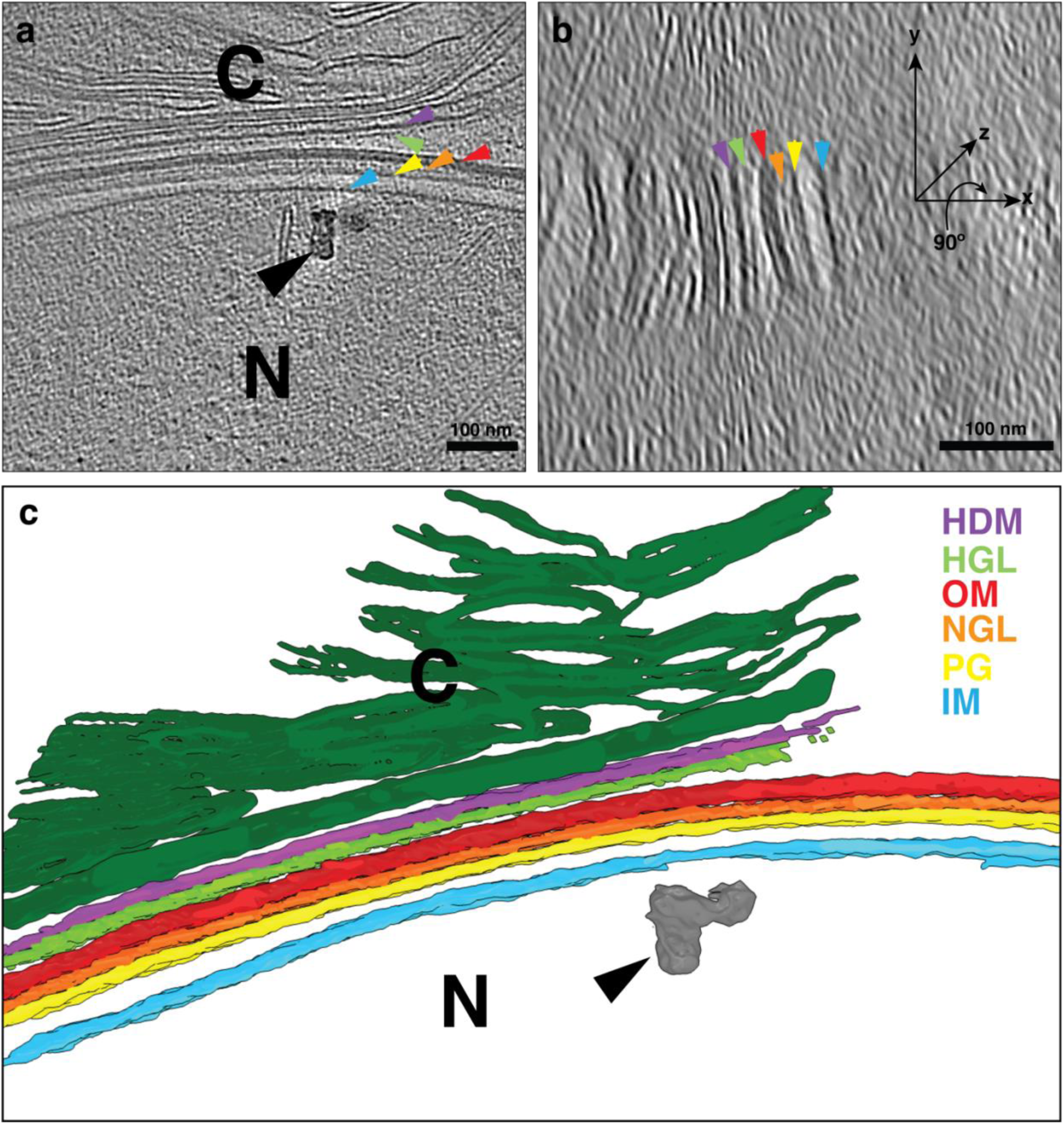
The composite envelope of the nitroplast. a, Slice through a cryo-electron tomogram of a morning nitroplast. **b,** Orthogonal cross-section (yz plane) of the same tomogram highlighting the six-layer envelope architecture: nitroplast inner membrane (IM), peptidoglycan (PG), nitroplast granular layer (NGL), nitroplast outer membrane (OM), host-derived membrane (HDM), and host granular layer (HGL). **c,** 3D segmentation of the tomogram in **a,b**. N, nitroplast; C, chloroplast. Arrow color code: IM (blue), PG (yellow), NGL (orange), OM (red), HGL (green), HDM (purple). Black arrow in **a** and **c** indicates an amorphous electron-dense body adjacent to the nitroplast IM.

In one cryo-tomogram, we observed three small (∼50 nm) spiky vesicles with dense central cores located in the space between the nitroplast OM and the host derived layers (HDM+HGL) (Fig. S7c). These observations point to an elaborated, multi-layered interface between the nitroplast and host cytoplasm that exceeds the complexity of known plastid envelopes.

### Nitroplast interfaces with multiple host organelles through distinct membrane contact sites

Consistent with FIB–SEM reconstructions, cryo-ET revealed, at higher resolution, extensive contact between the nitroplast and other host organelles (Fig. S8; Movies S8–S10), where spacing between the nitroplast and these other organelles ranged between 10-20 nm suggesting that these are genuine membrane contact sites (MCSs)^45^ (Fig. S9). When adjacent to mitochondria, an additional electron-dense layer was observed between the mitochondrial outer membrane and the nitroplast HDM (Fig. S8b,c), suggesting the presence of a specialized contact site or tethers. At nitroplast–chloroplast interface, ribosomes were observed tethered to the HDM, and in one instance a prominent spiral structure was anchored to the HDM at the interface between the two organelles (Fig. S8d,e, Movie S9). These features suggest localized translation or scaffolding at organelle contact sites. The nitroplast was also frequently adjacent (∼15 nm) to a previously uncharacterized mesh of branching membrane tubules (70–120 nm diameter) containing internal densities and encapsulated within a membrane and some tubules could be seen attached to this encapsulating membrane (Figs. S2&8 and Movie S10). Although no direct fusion or contact events were observed, the repeated proximity of this compartment may suggest a role in material trafficking or regulation. Collectively, these MCSs suggest a complex network of interactions between the nitroplast and other host cellular organelles.

### Diel remodeling of the nitroplast–host interface during nitrogen fixation

To determine whether nitroplast architecture changes across the light–dark cycle, we examined cells collected after 6 h of darkness (“night”) or 6 h of light (“day”). We collected 15 tomograms from 15-night cells and 8 tomograms from 7-day cells. Night nitroplasts closely resembled morning nitroplasts, retaining an intact HDM and HGL, a four-layer cyanobacterial envelope, and abundant triangular thylakoid-associated complexes (Fig. 4a–f, Movie S11). Subtomogram averaging of this triangular complex revealed a ∼20 nm-wide structure anchored to the membrane (Fig. 4f).

**Figure 4.**
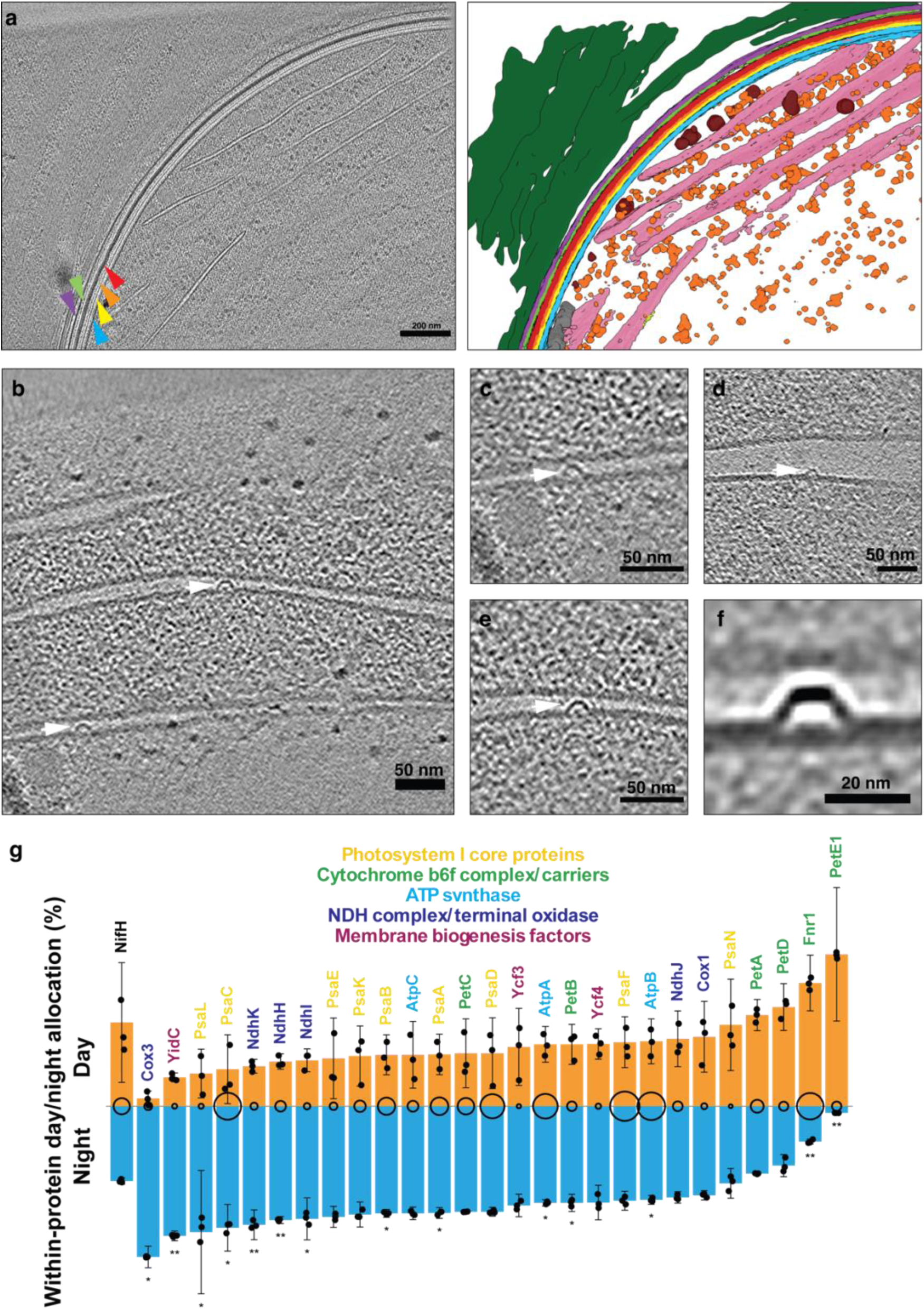
Structural features of the night nitroplast and diel regulation of a thylakoid-associated complex. a, Left: slice through a cryo-electron tomogram of a night nitroplast. Right: corresponding 3D segmentation. Envelope layers are indicated using the color code in Fig. 3. N, nitroplast; C, chloroplast. **b–e,** Tomographic slices highlighting triangular thylakoid-associated complexes (white arrows) in night nitroplasts. **f,** Central slice through a subtomogram average of the triangular complex (n = 10 particles). **g,** Comparative day/night allocation of thylakoid-associated proteins. Bars show normalized mean protein quantities (day + night = 100%); error bars indicate ±SD (n = 3 biological replicates, plotted individually as black dots). Circle size is proportional to overall mean abundance. Significance: *P* < 0.05, **P** < 0.01 (limma with Benjamini–Hochberg FDR correction). NifH is shown as a high-expression reference. Data replotted from ref. 5 and available in Data S1.

In striking contrast, day nitroplasts exhibited profound structural remodeling, and notably, the triangular thylakoid-associated complexes were absent from all day nitroplasts examined (Fig. S10). While the identity of this complex is unknown, proteomics analysis indicated that many prominent thylakoid-associated proteins exhibit diel changes in abundance, including many with increased abundance at night (Fig. 4g). Moreover, the HDM and HGL were frequently discontinuous or absent, exposing the nitroplast directly to the host cytoplasm and adjacent organelles (Fig. 5a,b, Movie S12). Concomitantly, we observed a significant flux in spiky vesicles dispersed throughout the cytoplasm near the nitroplast (>90 vesicles in day samples versus <10 in morning or night samples; Fig. S10). These spiky vesicles ranged from 30–130 nm in diameter and consistently contained a dense central granular core (∼25 nm; Fig. 5). Subtomogram averaging of vesicles with a diameter of ∼50 nm confirmed the uniform size and position of this core (Fig. 5g–k). The vesicle membrane closely resembles the nitroplast OM in thickness and density profile (Fig. S11), with the central granular core inside the vesicle similar to the GL present underneath the nitroplast OM (Fig. 5). These morphological similarities suggest that these vesicles originate from the nitroplast OM and GL. In addition, day nitroplasts were associated with smooth vesicles (50–350 nm) containing amorphous electron-dense material similar to that found within the nitroplast matrix (Fig. 5e,f; Fig. S12).

**Figure 5.**
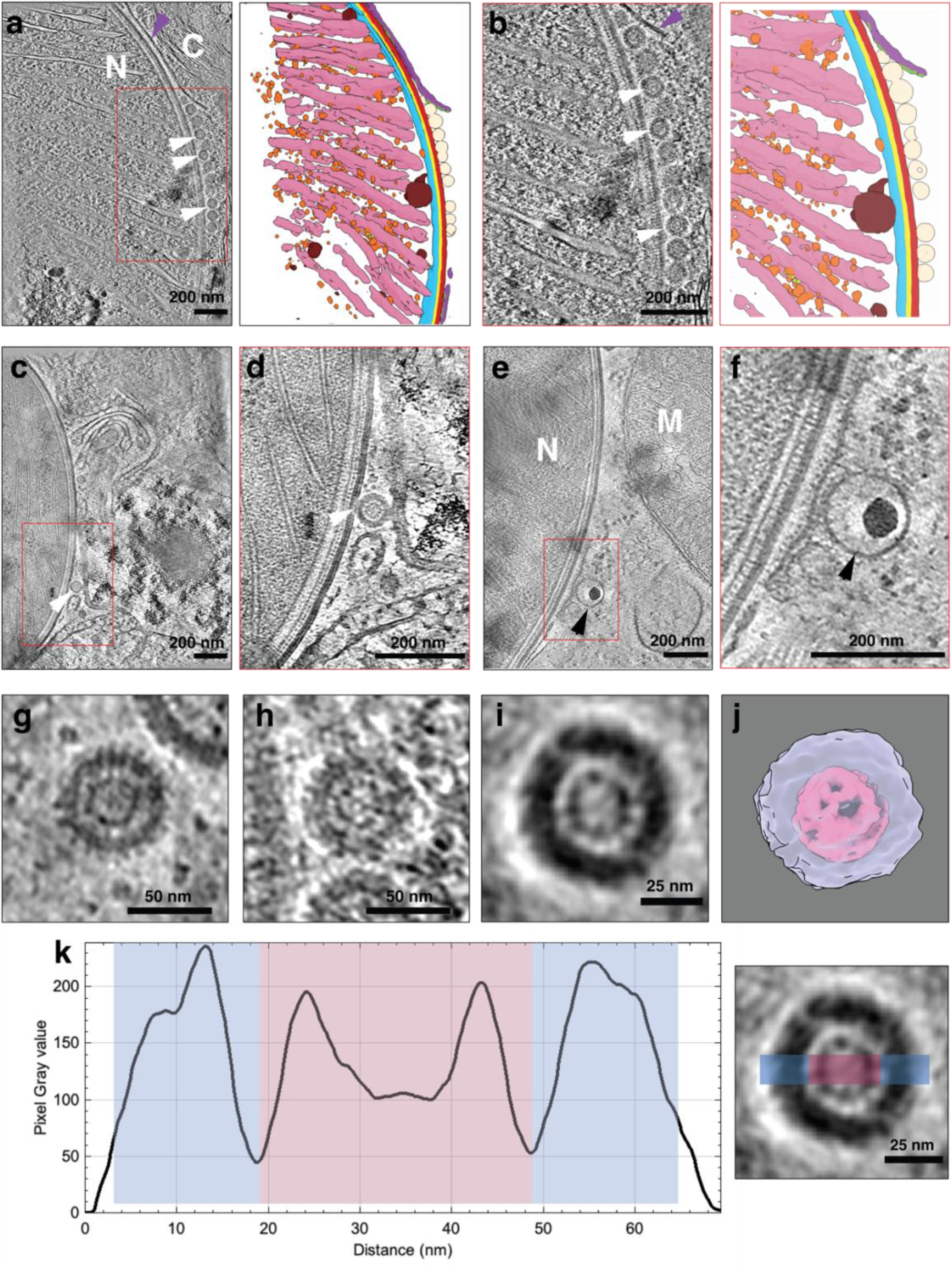
Daytime remodeling of the nitroplast–host interface and vesicle accumulation revealed by cryo-ET. a, Left: slice through a cryo-electron tomogram of a day nitroplast. Right: corresponding 3D segmentation highlighting discontinuities in the HDM (purple arrow) and the accumulation of spiky vesicles (white arrows). **b,** Enlarged view of the boxed region in **a**. **c,d,** Example of a spiky vesicle in direct contact with the nitroplast OM (white arrow in **c**); **d** shows an enlarged view the boxed region. **e,f,** Example of a smooth vesicle containing electron-dense cargo (black arrow in **e**); **f** shows an enlarged view of the boxed region. **g,h,** Additional examples of spiky vesicles. **i,** Subtomogram average of manually picked spiky vesicles (50–60 nm diameter) revealing a conserved central core. **j,** 3D visualization of the average shown in **i**. **k,** Left: average density profile across the region indicated in the right panel; peaks corresponding to the vesicle membrane are highlighted in light blue and peaks corresponding to the vesicle core are highlighted in pink.

Together, these observations define a diel structural cycle in which the nitroplast transitions from a membrane-isolated state at night to a vesicle-rich, cytoplasm-accessible state during daytime N_2_ fixation (Fig. 6).

**Figure 6.**
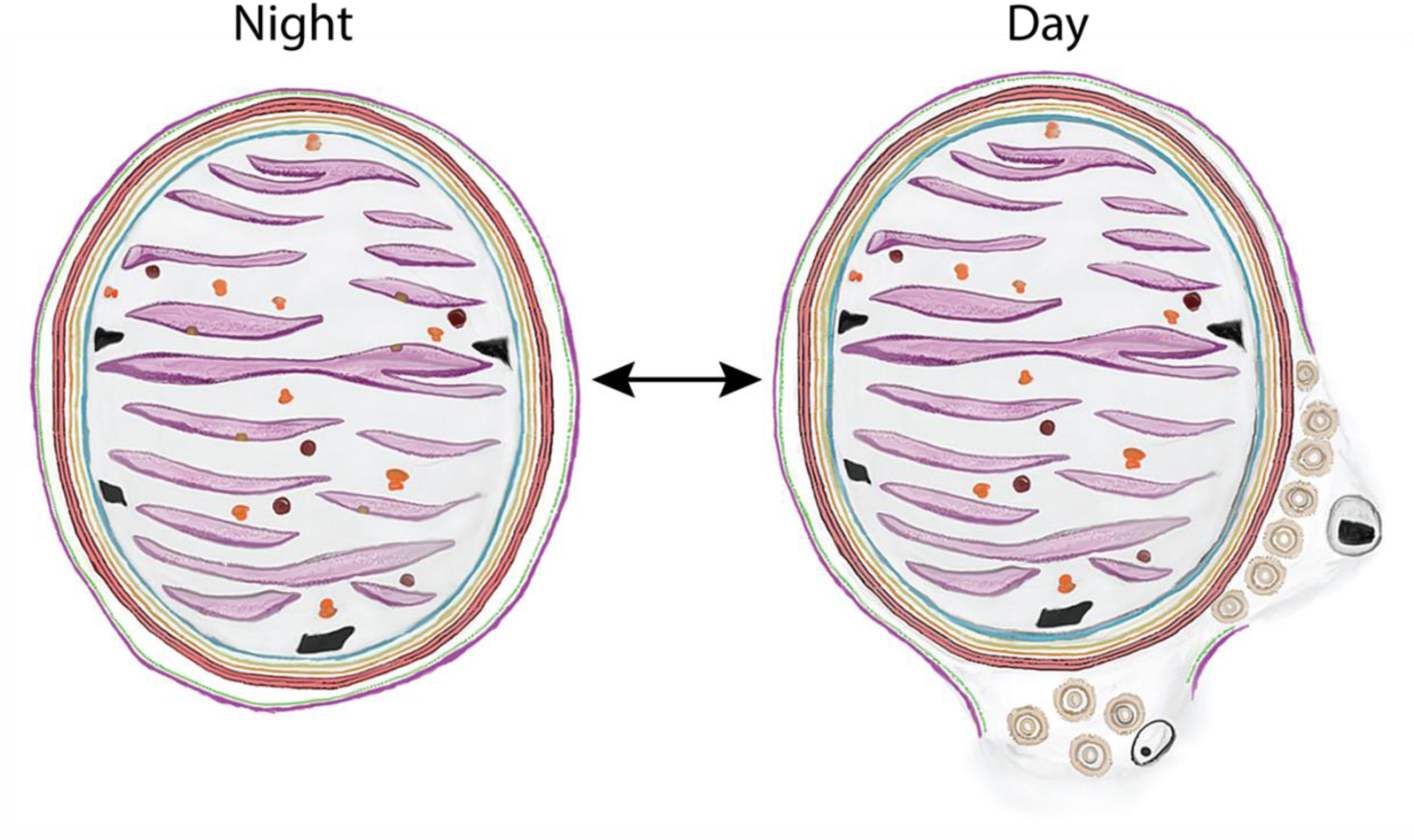
Model for diel remodeling of nitroplast architecture and host interface. Schematic summarizing major structural states across the diel cycle. At night, the nitroplast is enclosed by host-derived layers (HDM and HGL) and contains the triangular thylakoid-associated complex. During the day, host-derived layers become discontinuous and two vesicle populations accumulate at the nitroplast–host interface: spiky vesicles and smooth vesicles containing electron-dense cargo.

## Discussion

### Nitroplast integration preserves host architecture while enabling a costly metabolic function

The nitroplast, together with the chromatophores in *Paulinella*^46–48^, represent the most recently evolved eukaryotic organelles known to date, offering a rare opportunity to examine the early stages of organellogenesis. By combining volume electron microscopy with *in situ* cryo-ET from cultured and freshly collected cells from the ocean, we provide a comprehensive view of how a N_2_ fixing endosymbiotic organelle is structurally integrated, regulated, and remodeled within a eukaryotic cell.

Our comparative analyses with closely related microalgae lacking a nitroplast indicate that nitroplast acquisition, despite occupying 10% of the cell, did not affect host organelle scaling or cellular architecture, despite the high energetic cost of N_2_ fixation^49^. The proportional volume of chloroplasts and mitochondria remains conserved relative to closely related haptophytes lacking a nitroplast. This is unexpected, as N_2_ fixation is energetically expensive, requiring 16 ATP per N_2_ reduced. Nonetheless, part of this energy could be supplied by the nitroplast itself given that it has retained photosystem I^41^. This suggests that the host cell accommodates N_2_ fixation not by expanding existing energy-producing organelles, but through efficient coupling of nitroplast metabolism to existing cellular structures, likely including retained photosystem I activity within the nitroplast itself. However, additional compensatory mechanisms involving, for example, mitochondrial cristae architecture, may further contribute to meeting the energetic demands imposed by N_2_ fixation. Together, these findings indicate that nitroplast integration preserves the fundamental architecture of the host cell, despite the addition of a large, metabolically demanding organelle.

Notably, the conserved chloroplast-to-nitroplast volume ratio across calcified and non-calcified life stages suggests a tightly regulated balance between host carbon fixation and nitroplast nitrogen fixation capacity. This invariant scaling is particularly striking given the substantial differences in absolute cell size, morphology, and nitroplast abundance between these stages, and argues against a passive association. Instead, it is consistent with quantitative coupling between photosynthetic output and the energetic and reductive demands of N_2_ fixation. Recent work further supports that the size ratio between UCYN–A and the chloroplasts of their respective hosts is strikingly conserved across sublineages and species, including not only the nitroplast (i.e., UCYN–A2) but also smaller UCYN–A–derived endosymbionts such as UCYN–A1 and UCYN–A3 sublineages, thus indicating that metabolic interdependence likely imposes strict constraints on organelle volume^14^. Given that N₂ fixation is highly ATP-intensive and depends on carbon supplied by the host, maintaining a fixed proportional relationship between these compartments may ensure coordinated metabolic flux and redox balance. The reduced relative investment in nitroplast volume observed in calcified cells, together with their larger size, may reflect differences in metabolic state or nitrogen demand across life stages or environmental conditions. More broadly, this conserved scaling supports the view that the nitroplast is functionally integrated into host cellular physiology and regulated in concert with other core metabolic organelles.

In oligotrophic ocean waters, where fixed (bioavailable) nitrogen (nitrate, nitrite and ammonium) occurs at nanomolar concentrations, importing and reducing available nitrogen species also imposes significant energetic costs. Uptake of fixed nitrogen species against steep concentration gradients requires active transporters and ATP-driven assimilation pathways, including nitrate and nitrite reductases, and associated electron-transfer proteins^50,51^. In contrast, direct intracellular N_2_ fixation by the nitroplast circumvents these transport and reduction steps, especially if the process is tightly coupled to photosynthetically derived energy. Importantly, *B. bigelowii* appears to have lost the capacity to use nitrate^31^, remains actively N_2_-fixing even in nitrogen-replete coastal waters^52^, and even large nitrate additions do not suppress N_2_ fixation in culture^13^, consistent with the host relying on nitroplast-derived fixed N for a large fraction of its demand. Consequently, we expect the physiology and ultrastructure described here to be representative of an actively N_2_-fixing organelle supplying the bulk of the host cell’s fixed-nitrogen requirement^32^. Additionally, decreased investment in nitrogen importers could free membrane surface area and reallocate cellular resources (energy, amino acids, and cofactors) to uptake of other growth-limiting nutrients. Thus, when efficiently integrated into host metabolism, the energetic burden of N_2_ fixation may be comparable to, rather than greater than, the cost of scavenging and assimilating nitrogen in oligotrophic environments.

### The nitroplast is physically integrated into the host through membrane contact sites and a hybrid envelope architecture

At the cellular level, the nitroplast forms multiple membrane contact sites (MCSs) with other host organelles. MCSs are discrete regions where two organelles are closely apposed—typically within ∼30–50 nm—often bridged by tethering factors, and they have emerged as key hubs for coordinating organelle function and inter-organelle exchange, including in mitochondria^45,53^. In our cryo-ET data, the appearance of an additional layer at close (∼ 5 nm) nitroplast HDM–mitochondrion appositions is consistent with a specialized contact site that could mediate tethering and coordinate metabolic coupling between these compartments in the microalga *B. bigelowii*. In particular, the proximity between the nitroplast and mitochondria may suggest a potential role for mitochondrial metabolism in supporting N_2_ fixation. Analogous structures may underlie the recurrent nitroplast–chloroplast interfaces, where we observe putative tethers and, in one instance, a prominent spiral density in direct contact with the HDM. This nitroplast’s proximity to chloroplasts implies potential mechanisms that buffer or divert photosynthetically evolved O_2_, enabling close association that may facilitate metabolite exchange. Notably, the nitroplast also approaches the branched tubular meshwork to within ∼15 nm, suggesting a direct physical interaction between these compartments as well. Together, these observations point to a structured network of contacts linking the nitroplast to multiple host organelles, which may help integrate this recently acquired, oxygen-sensitive compartment into cellular physiology by enabling regulated exchange and local coordination of metabolism.

At the structural level, the nitroplast exhibits a hybrid envelope architecture. It retains a reinforced cyanobacterial envelope including PG and a thick OM, while acquiring additional host-derived membrane layers that are dynamically regulated. This composite architecture distinguishes the nitroplast from both free-living cyanobacteria and canonical plastids, which have largely lost PG and bacterial envelope features. In primary plastids of green algae and plants, the PG layer is usually lost or reduced and envelopes are reduced to two membranes (former IM and OM), whereas complex plastids of secondary/tertiary endosymbiosis add one to two host-derived membranes but still lack PG^54,55^. By contrast, marine cyanobacteria commonly retain reinforced envelopes that either exhibit an external layer beyond the OM^56^, a conspicuously thick OM^57^, or an additional fourth layer located between the OM and the PG^58^. Among marine diazotrophs, the non-heterocyst-forming filamentous *Trichodesmium* retains a Gram-negative OM–peptidoglycan–IM envelope^59^, whereas heterocyst-forming symbionts (e.g., *Richelia* in diatom–diazotroph associations) include thick walls and glycolipid outer layers that lower O₂ permeability^60,61^. The persistence of bacterial structural elements likely reflects the recent evolutionary origin of the nitroplast and the need to protect oxygen-sensitive nitrogenase, while more ancient cyanobacteria-derived organelles have typically lost the PG layer^62^.

An additional feature that may reflect the recent evolutionary origin of the nitroplast is the presence of tubular mesh structures in *B. bigelowii* cells that are occasionally adjacent to the nitroplast observed in both imaging methods on several cells. While the nature of these structures remains unknown, it bears a striking morphological similarity, at the resolution of our cryo-tomograms, to a mesh of membrane tubes present at the host-microbe interface of the symbiosis between plant root cells and the arbuscular mycorrhizal fungi that is presumably involved in materials exchange between the two symbionts^63–65^. It is possible that a similar system was used to communicate between UCYN-A and the host in the symbiosis stage and that it has been retained as UCYN-A continues to evolve from a free-living bacterium to a cellular organelle.

### Diel remodeling reveals regulated exchange across the nitroplast–host interface

Perhaps the most striking observation is the diel remodeling of the nitroplast–host interface. During the day, when N_2_ fixation occurs, the HDM barrier becomes discontinuous and the organelle is surrounded by abundant vesicles. At night, the nitroplast is enclosed by host membranes. This temporal gating suggests a regulatory strategy in which the host alternates between isolating the nitroplast to limit exchange and opening access to cytosolic metabolites when N_2_ fixation is active. More generally, these observations could suggest that early organellogenesis may proceed through temporally gated modulation of membrane barriers, rather than immediate establishment of static organelle boundaries.

The spiky vesicles that accumulate during the day are particularly intriguing. Their morphological similarity to the nitroplast OM and the presence of a granular core at their center suggest a specialized trafficking function originating from the nitroplast. If these vesicles indeed originate from the nitroplast, the granular core inside them could derive from the GL of the nitroplast. This is consistent with the observation of these spiky vesicles, at a much lower frequency, in the space between the nitroplast OM and the HDM in the morning nitroplast when N_2_ fixation is presumably beginning. While the molecular cargo and directionality of these vesicles remain to be determined, their temporal specificity suggest a regulated exchange pathway.

Extracellular vesicle production is increasingly recognized as a common feature of cyanobacteria, including dominant marine lineages (e.g., *Prochlorococcus* and *Synechococcus*), and in Gram-negative bacteria these structures most often correspond to outer-membrane vesicles enriched in bacterial envelope components and carrying characteristic lipids and proteins^66,67^. Vesiculation has also been reported in model freshwater cyanobacteria^68^, supporting outer-membrane vesicle production as an intrinsic feature of cyanobacterial envelope biology. Notably, abundant vesicle-like structures have been reported at the envelopes of intracellular N_2_-fixing symbionts, including periplasmic vesicles and apparent outer-membrane evaginations/extrusions, suggesting a role in facilitating metabolite exchange and tight metabolic coupling in N_2_-fixing symbioses^69^. Together, these precedents support the possibility that the vesicles observed at the nitroplast periphery originate from the nitroplast envelope rather than from host membranes; however, definitive assignment will require further evidence.

The smooth vesicles that contain amorphous electron-dense material may represent an additional trafficking pathway. Their resemblance to dense bodies within the nitroplast matrix raises the possibility that they function as iron reservoirs or transport intermediates, supplying iron-rich cofactors required for nitrogenase, ferredoxin, and photosystem I^13^. An iron reservoir is crucial for the function of the nitroplast which lacks the ability to obtain iron directly from the environment due to its physical location inside the *B. bigelowii* cell. Such a transport mechanism would parallel phytotransferrin-mediated iron trafficking to chloroplasts in marine phytoplankton^70,71^. Future experiments utilizing techniques such as electron energy loss spectroscopy might shed light on the content of these vesicles.

Finally, the unidentified triangular complexes embedded in nitroplast thylakoids exhibit diel regulation and structural similarity to membrane-associated assemblies in both bacteria^72,73^ and other organelles (e.g., chloroplast^74^ and mitochondria^75^). Consistent with this structural remodeling, our nitroplast proteomics shows pronounced diel regulation across multiple thylakoid-associated proteins, including several highly abundant PSI/electron-transfer and ATP synthase components that are relatively enriched at night, supporting the idea that nitroplast thylakoids are functionally reconfigured over the diel cycle and potentially linking the triangular complex to these nighttime thylakoid states. Their disappearance during the day could suggest a role decoupled from active N_2_ fixation. Together with the membrane remodeling we observe at the organelle periphery, these changes point to a broader reorganization of nitroplast membrane systems across the diel cycle.

### Limitations of structural inference and implications for early organellogenesis

While our analyses provide direct insight into the structural remodeling and regulation of the nitroplast, they also define clear boundaries of what can be inferred from static architectural data alone. The temporal coincidence of membrane remodeling with the known window of N_2_ fixation provides strong circumstantial evidence for regulatory coupling, although additional light-dependent processes may also contribute to this remodeling. Our data therefore establish a structural framework within which regulated host–nitroplast exchange can occur, rather than identifying the specific molecular substrates involved. Similarly, the molecular cargo and directionality of the two vesicle populations observed at the nitroplast interface during the day remain to be determined.

In summary, our study supports a model in which early organellogenesis proceeds through dynamic, temporally regulated membrane remodeling rather than static encapsulation. Ultimately, the evolutionary trajectories of mitochondria and plastids rendered them permanently accessible to the host cytoplasm through the reduction of their bacterial envelopes and the acquisition of dedicated molecular transport machineries^9,62,76–78^. If an intermediate stage with restricted cytoplasmic access existed during the evolution of these organelles, it was likely transient and subsequently vanished as envelope reduction and specialized transport systems emerged, leaving little detectable trace in extant lineages. The nitroplast therefore exemplifies how a eukaryotic cell can integrate a new, energy-intensive metabolic function by balancing bacterial structural inheritance with host-controlled elements and exchange. While the nitroplast is unique, the regulatory strategy we uncovered, including dynamic control of organelle accessibility, may represent a broadly applicable solution for integrating metabolically specialized compartments. Beyond its evolutionary significance, this system provides a natural blueprint for efforts to engineer N_2_-fixing compartments in eukaryotic cells, linking fundamental cell biology to future biotechnological applications^79^.

## Acknowledgements

This work was supported by the National Institute of General Medical Sciences (R35GM157116 to M.K.) and the Searle Scholars Program (to M.K.), the Simons Foundation (#824082 and # SFI-LS-Project-00010362 to J. P. Z), and the Gordon and Betty Moore Foundation (AtlaSymbio project: https://doi.org/10.37807/GBMF11532 to J. D.). JD and TD were supported by the ERC consolidator grant SymbiOCEAN (101088661). Ethan Li was supported by the QUAD Undergraduate Research Scholars Program from the University of Chicago. F.M.C.-C. was supported by the Spanish Ministry of Science, Innovation and Universities (MICINN) through the grant COMMUNAS (PID2023-152110NA-I00), a ‘Ramón y Cajal’ fellowship (RYC2021-032949-I), and the ‘Severo Ochoa Centre of Excellence’ accreditation (CEX2024-001494-S) funded by AEI 10.13039/501100011033. We acknowledge the technical support provided by The Advanced Electron Microscopy Facility at the University of Chicago, the Biomolecular Cryo-Electron Microscopy Facility at the Department of Chemistry and Biochemistry of the University of California, Santa Cruz (RRID:SCR_021755; NIH S10 High-End Instrumentation program S10OD02509), and the Electron Microscopy Core facility at EMBL, Heidelberg. We thank Benoit Gallet, Guy Schoehn, Christine Moriscot, Emma Dusacq and the electron microscope facility at IBS, which is supported by the Rhône-Alpes Region, the Fondation Recherche Medicale (FRM), the fonds FEDER, the Center National de la Recherche Scientifique (CNRS), the CEA, the University of Grenoble, EMBL, and the GIS Infrastructures en Biologie Sante et Agronomie (IBISA). We also thank the TREC expedition consortium and the Advanced Mobile Laboratory of the European Molecular Biology Laboratory (EMBL), in particular Niko Leish and Paulina Cherek. For the support with environmental sampling, we thank the Traversing European Coastlines (TREC) Consortium, the TREC core partners EMBL, the Tara Ocean Foundation, the Tara Europa Consortium and the European Marine Biological Resource Centre (EMBRC) for their commitment to making the TREC expedition possible, and especially the Kristineberg Centre for Marine Research and Innovation where samples were collected for some results shown here.

We are grateful to Benjamin Glick (University of Chicago) and Samuel H Light (University of Chicago) for critically reading the manuscript and Douglas C. Rees (Caltech) and Matthew Swulius (Pennsylvania State University) for insightful discussions.

## Materials and Methods

### Strains and growth conditions

*B. bigelowii* strain FR-2^13^ was maintained in batch culture at 18 °C and 60 μmol photons m^-2^ s^-1^ under a 12 h:12 h light–dark cycle, and contained in 250 mL polycarbonate bottles. Cultures were grown in sterilized natural Monterey Bay seawater (36°56’53.5”N 122°03’55.2”W) amended with f/2 macronutrients (880 μM NO_3_ ^-^, 36 μM PO ^3-^), vitamins (296 nM thiamine, 2.05 nM biotin, 0.37 nM cobalamin) and trace metals (200 nM Fe, 100 nM Zn, 48 nM Mn, 40 nM Co, 40 nM Cu, 10 nM Se and 100 nM Ni, supplied with 100 μM EDTA as a chelator). For the FIB-SEM experiment, *B. bigelowii* strain FR-21 was maintained at 16°C and 60 μmol photons m^-2^ s^-1^ under a 10 h:14 h light-dark cycle, and contained in 250 mL cell culture flasks. Cultures were grown in sterilized natural Blanes Bay seawater (41°40′N, 2°48′E) amended with Guillard F/2 media^80^, and supplemented with filtered gelidium jelly extract (GJE) as described in reference^31^. Prior to use, seawater was adjusted to pH 8.1 and salinity 34 ppt, autoclaved, and then amended with nutrients, vitamins and trace metals as described above. Cultures were maintained in exponential phase by regular dilution into fresh medium prepared in the same manner, and cell concentrations were monitored in culture aliquots using a BD Accuri^TM^ C6 Plus flow cytometer (BD Biosciences, Franklin Lakes, NJ, USA).

### Sample preparation and FIB-SEM imaging of cultured cells

Samples of *B. bigelowii* strain FR-21 cultures in exponential phase (cell density ∼100,000 cells mL^-1^) were collected 6 hours after lights on, concentrated in an Amicon® Ultra-4 (Merck, UFC810096) by centrifugation at 2300 g for four minutes at 21°C. The concentrated *B. bigelowii* strain FR-21 culture was cryofixed using High Pressure Freezing (HPM100, Leica Microsystems, Austria) at a pressure of 210 MPa at −196°C in the Advanced Mobile Laboratory (AML) as described in references^38,81^. This was followed by freeze-substitution (EM ASF2, Leica Microsystems, Austria), where vitrified ice was replaced with dried acetone and 2% osmium tetroxide, and then embedded in Epon for FIB-SEM. More details of the sample preparation protocols are provided in protocols.io^82^. The resin blocks were then stored in dry conditions prior to imaging.

Focused Ion beam Scanning Electron Microscopy (FIB-SEM) tomography was performed with a Zeiss CrossBeam 550 microscope with the Atlas 5 software (Zeiss, Germany). The resin block containing the cells was fixed on a stub with silver paste and surface abraded with a diamond knife in a microtome to obtain a flat and clean surface. Samples were then metallized with 8 nm of platinum to avoid charging during observations. Inside the FIB-SEM, a second platinum layer (2 µm) was deposited locally on the analyzed area to mitigate possible curtaining artifacts. The sample was then abraded slice by slice with the Ga+ ion beam (typically with a current of 700 pA at 30 kV). Each exposed surface was imaged by SEM at 1.5 kV and with a current of 1.5 nA using the in-column EsB backscatter detector. Similar milling and imaging modes were used for all samples. Automatic focus and astigmatism correction were performed during image acquisition, typically at approximately hourly intervals. For each slice, a thickness of 8 nm was removed and SEM images were recorded with a corresponding pixel size of 8 nm in order to obtain an isotropic voxel size. Entire volumes were imaged with ∼500-800 frames, depending on the cell type and volume. The first steps of image processing were performed using Fiji software for registration (adapted StackReg plugin), and noise reduction (3D mean function of the 3D suite plugin^83^). Registered electron microscopy data have been deposited in the Electron Microscopy Public Image Archive (EMPIAR), accession code EMPIAR-XXX.

Semi-automatic and manual segmentation using 3D Slicer^84^ was used to obtain volumes of organelles. Plastids, nitroplast, mitochondria and nuclei of the microalga were segmented to reconstruct the cellular organization and quantify volumes (mm3). Each segment (e.g. nitroplast, plastid, mitochondria) was colored using paint tools, adjusting the threshold range of the pixel values (intensity) of the images. Different views and slices were generated for the model using the EasyClip module. Volume measurements were then calculated using the Segmentation Statistics module. Paraview^85^ was also used to edit the 3D models and generate images and videos.

### Environmental sampling and sample preparation for electron microscopy

Natural phytoplankton communities were sampled with a 10 µm net near the Kristineberg Centre for Marine Research and Innovation during the TREC expedition (https://www.embl.org/about/info/trec/) and subsequently size-fractionated using a 40 µm sieve. Samples were processed in the Advanced Mobile Laboratory (AML)^86^ deployed on site. The sample was concentrated onto a 1.2 µm mixed cellulose ester membrane using a manual vacuum filtration unit, and allowed to sediment prior to high-pressure freezing (EM-ICE, Leica Microsystems). A 1.2 µl aliquot of the concentrated mix was transferred into a type A carrier (gold-coated copper), 3mm wide and 200 µm deep, and sandwiched with a type B aluminum carrier (Wohlwend).

Sample was freeze substituted (EM-AFS2, Leica Microsystems), following protocols previously described^40,87^. The sample was incubated in 0.1% uranyl acetate dissolved in dry acetone at −90°C for 72 hours. The temperature was then gradually increased to −45°C at 2°C per hour, followed by incubation at −45°C for 10 hours. Resin infiltration was carried out with progressively increasing concentrations (10%, 25%, 50%, 75%, and 100%) of Lowicryl HM20 while raising the temperature to −25°C. After three exchanges with 100% Lowicryl resin, each lasting 10 hours, polymerization was carried out using UV light at −25°C for 48 hours. Finally, the temperature was increased to 20°C, and UV polymerization was continued for an additional 48 hours.

A 3D confocal map of the resin block was acquired using a Zeiss LSM 780 NLO microscope. A 4X4 tiled scan was acquired using the 25X/0.8 NA multi-immersion objective. Cells of interest were identified and branded with the 2-photon laser set to 800 nm at 10% power. The block was then mounted onto a SEM stub. Silver paint was applied around the sample area, and the stub was sputtered with gold at 30 mA for 180s (Q150RS, Quorum).

### FIB-SEM imaging of environmental cells

3 cells in coccolith stage were acquired at an isotropic 10 nm voxel size at the Zeiss Crossbeam 540. Milling was done at 30 kV, 3 nA and SEM imaging at 1.5 kV accelerating voltage and current of 700 pA with an ESB detector (ESB Grid 1110V). The final dwell time for acquisition was 11-12 µs. Raw datasets were cropped and aligned using the automated Alignment to Median Smoothed Template (AMST) workflow described by reference^88^ for cells 1 and 2, and the Fiji plugin Linear Stack Alignment with SIFT for cell 3. Organelles were segmented from the FIB-SEM stack using Amira (Thermo Fisher Scientific) for visualization and morphometric analysis.

### Cryo-electron tomography sample preparation

*B. bigelowii* cultures were grown to exponential phase (∼100,000 cells mL^-1^) at which point subsamples were taken at 3 time points (1 hour after lights on, 6 hours after lights on, and 6 hours after lights off). Multiple grids were prepared from each time point by applying 4 µL of cells to glow-discharged Quantifoil R2/2, 200-mesh grids. The grids were rapidly vitrified in liquid ethane using a Leica GP2 cryo-plunger operated at 85% humidity and 18 °C, with a blotting time of 2 s in sensor blotting mode. Initial grid screening was conducted at the UCSC Biomolecular cryo-EM Facility on a Thermo Fisher Scientific Glacios 200 kV microscope equipped with a Gatan K2 Summit detector. Pre-clipped grids mounted CryoFIB AutoGrid, suitable for subsequent cryo-FIB milling, were imaged at 46× magnification with a 1 s exposure. The best-performing grids were shipped to the University of Chicago cryo-EM Facility for further lamella preparation and high-resolution data acquisition.

### Cryo-focused ion beam milling

For cryo-FIB milling, grids were clipped in CryoFIB AutoGrid Rings (Thermo Fisher Scientific) with a cutout to allow for shallower milling angles. Milling was performed on an Aquilos 2 cryo-DualBeam instrument (Thermo Fisher Scientific). Samples were sputter-coated in-column with platinum for 20 s at 10 Pa and 20 mA, and then they were coated with a layer of organometallic platinum for 30 s using the gas injection system within the instrument. Targets were identified and marked using Maps v. 3.33 software and milled using AutoTEM v. 2.4 software (Thermo Fisher Scientific). The automated protocol included milling of micro-expansion joints at 0.3 nA, followed by rough, medium, and fine milling at 0.3, 0.1 and 0.05 nA, respectively, followed by two polishing steps at 30 and 10 pA. The result was a lamella with a target thickness of 200 nm. Milled samples were stored under liquid nitrogen until cryo-ET imaging was performed.

### Cryo-ET data collection and image processing

For cryo-ET imaging, data were collected on a Titan Krios G3i at 300 kV (Thermo Fisher Scientific) with a Gatan K3 direct detection camera in CDS mode with the initial dose rate target on the detector between 7.5-8.5 electrons per pixel per second and with the Gatan BioQuantum-K3 energy slit width set to 20 eV. Tomography 5 software (Thermo Fisher Scientific) was used for collection. The tilt series was acquired with 3° steps in a bidirectional collection scheme^89^ beginning with a lamella pre-tilt of ±9° and extending to ±54° or 60°. The total dose for each tilt series was 120 e-/Å^2^. Tilts were acquired with a pixel size of 4.45, 3.36, or 1.68 Å and with a defocus range of 5 to 8 µm. Tilt series data were reconstructed using the IMOD software package^90^. 3D segmentation was done using Dragonfly software (https://www.theobjects.com/dragonfly/index.html).

The subtomogram averaging of the hat-like structure was done using the PEET program^91^ with two-fold symmetrization applied along the particle *Y* axis. The number of particles averaged was 10 particles. The subtomogram averaging of the vesicles was done using Dynamo^92,93^ (1.1.555 Matlab R_2025a version) with the application and all the corresponding scripts being run on Matlab R_2025a. Sixteen nitroplast vesicles were manually picked and averaged.

### Statistical Analysis

All statistical analyses and data visualizations were performed using R, running in RStudio. The ggplot2 package was used for all data visualization. For all statistical tests, a significance level of α = 0.05 was used.

Group Comparisons: To compare morphological features between day and night samples, Welch’s Two Sample t-tests were employed. These were chosen as it does not assume equal variance between the two groups. Comparisons were made for the counts of globules, vesicles, and hat-like structures. In addition, comparisons were made for the diameters of globules and vesicles.

Initial exploratory data analysis of vesicle diameters revealed a distribution with extreme values, and thus Welch’s t-test for vesicle diameters was conducted on the (i) complete dataset and (ii) a filtered dataset with outliers removed. Outliers were formally defined as any data point falling outside the bounds of Q_1_ – 1.5 x IQR and Q_3_ + 1.5 x IQR (where Q_1_ and Q3 represent the first and third quartiles, and IQR is the interquartile range).

Correlation Analysis: To determine if a linear relationship existed between the abundance and size of specific structures to the day or night state, Pearson’s product-moment correlation was calculated.

### Density Profile Plots

To analyze the interlayer spacing of the nitroplast membranes, membrane density profile plots were generated using Fiji (ImageJ). First, representative 2D slices showing clear membrane organization were extracted from the reconstructed tomogram files. Within ImageJ, a rectangular Region of Interest was drawn perpendicular to a well-aligned segment of the stacked membranes. This processing pipeline was then applied: grayscale inversion, noise reduction by applying a Gaussian blur filter, and contrast enhancement. The spatial scale was calibrated using the pixel size (nm/pixel) from the original tomogram metadata. Finally, the native “Plot Profile” was used to generate a 1D density plot that shows intensity peaks corresponding to the position of each layer.

### Nitroplast day-night proteomic analysis

Proteomics data were taken from the published reference ^13^ full protein-group intensity table provided in the accompanying Dryad repository (ADK1075_ProteinQuantifications.csv, https://doi.org/10.5061/dryad.2z34tmptf)^94^. We used only isolated UCYN-A samples collected during the day (n=3) and night (n=3). For visualization, protein-group quantity values were converted to within-sample relative abundances by dividing each protein’s quantity by the summed quantity across all protein groups in that sample. Day and night mean relative abundances were then calculated across replicates and within-protein normalized such that Day + Night = 1 (reported as %); for display (Fig 4g), night allocation is plotted as negative and day allocation as positive to facilitate mirrored comparison. Error bars denote ±1 SD across replicate relative abundances, propagated through the within-protein normalization. For statistical testing of day-night differences, we applied limma^95^ to log2-transformed protein-group quantities after between-sample median centering. P values were adjusted using the Benjamini–Hochberg procedure and significance was assessed at FDR < 0.05. Full analysis of day-night differential protein expression is available in Dataset S1.

## Movie legends

**Movie S1** FIB–SEM–based 3D reconstruction of a cultured, motile *B. bigelowii* cell showing chloroplasts (green), mitochondria (red), vacuole (purple), nucleus (blue), and nitroplast (yellow).

**Movie S2** FIB–SEM–based 3D reconstruction of a cultured, motile *B. bigelowii* cell highlighting mitochondria (red), vacuole (purple), nucleus (blue), and nitroplast (yellow).

**Movie S3** FIB–SEM image stack through a cultured, motile *B. bigelowii* cell illustrating the ultrastructural organization and spatial relationships of major organelles.

**Movie S4** FIB–SEM image stack of a calcified, non-motile *B. bigelowii* cell isolated from natural phytoplankton, followed by a 3D segmentation highlighting chloroplasts (green), nitroplast (yellow), and nucleus (blue).

**Movie S5** Cryo-electron tomogram of a cryo-FIB–milled morning nitroplast illustrating its internal thylakoid organization and overall morphology.

**Movie S6** Cryo-electron tomogram of a cryo-FIB–milled morning nitroplast illustrating its internal thylakoid organization and overall morphology.

**Movie S7** Cryo-electron tomogram of a cryo-FIB–milled morning nitroplast emphasizing the composite envelope architecture. Arrow color code: IM (blue), PG (yellow), NGL (orange), OM (red), HGL (green), HDM (purple).

**Movie S8** Cryo-electron tomogram of a cryo-FIB–milled morning nitroplast (N) in proximity to other organelles, including chloroplast (C), mitochondrion (M), vacuole (LV), and a tubular meshwork (T).

**Movie S9** Cryo-electron tomogram of a cryo-FIB–milled morning nitroplast (N) adjacent to a chloroplast (C), highlighting a spiral structure (boxed) anchored to the nitroplast HDM at the organelle interface.

**Movie S10** Cryo-electron tomogram of a cryo-FIB–milled morning nitroplast (N) positioned near an unidentified membrane-bound compartment containing a mesh of tubules (T).

**Movie S11** Cryo-electron tomogram of a cryo-FIB–milled night nitroplast adjacent to a chloroplast, followed by 3D segmentation highlighting the six-layer composite envelope and other internal structural features.

**Movie S12** Cryo-electron tomogram of a cryo-FIB–milled day nitroplast showing discontinuities in the HDM and accumulation of spiky vesicles at the nitroplast–host interface and other internal structural features.

**Data S1 Nitroplast day/night proteomics dataset.**

This dataset includes proteomics quantifications, limma differential-abundance results and per-replicate day/night allocation values used to generate Figure 4g.

## References

1. Field, C.B., Behrenfeld, M.J., Randerson, J.T., and Falkowski, P. (1998). Primary Production of the Biosphere: Integrating Terrestrial and Oceanic Components. Science 281, 237–240. 10.1126/science.281.5374.237.

2. Zehr, J.P. (2011). Nitrogen fixation by marine cyanobacteria. Trends in Microbiology 19, 162–173. 10.1016/j.tim.2010.12.004.

3. Eberhard, S., Finazzi, G., and Wollman, F.-A. (2008). The Dynamics of Photosynthesis. Annu. Rev. Genet. 42, 463–515. 10.1146/annurev.genet.42.110807.091452.

4. Bennett, C.F., Latorre-Muro, P., and Puigserver, P. (2022). Mechanisms of mitochondrial respiratory adaptation. Nat Rev Mol Cell Biol 23, 817–835. 10.1038/s41580-022-00506-6.

5. Koonin, E.V. (2010). The origin and early evolution of eukaryotes in the light of phylogenomics. Genome Biology 11, 209. 10.1186/gb-2010-11-5-209.

6. Archibald, J.M. (2009). The Puzzle of Plastid Evolution. Current Biology 19, R81–R88. 10.1016/j.cub.2008.11.067.

7. Elias, M., and Archibald, J.M. (2009). Sizing up the genomic footprint of endosymbiosis. BioEssays 31, 1273–1279. 10.1002/bies.200900117.

8. Archibald, J.M. (2015). Endosymbiosis and Eukaryotic Cell Evolution. Current Biology 25, R911–R921. 10.1016/j.cub.2015.07.055.

9. Keeling, P.J. (2010). The endosymbiotic origin, diversification and fate of plastids. Phil. Trans. R. Soc. B 365, 729–748. 10.1098/rstb.2009.0103.

10. Postgate, J.R. (1982). The fundamentals of nitrogen fixation (Cambridge University Press).

11. Grujcic, V., Mehrshad, M., Vigil-Stenman, T., Lundin, D., and Foster, R.A. (2025). Stepwise genome evolution from a facultative symbiont to an endosymbiont in the N2-fixing diatom-Richelia symbioses. Current Biology 35, 4479–4493.e3. 10.1016/j.cub.2025.08.003.

12. Fiore, C.L., Jarett, J.K., Olson, N.D., and Lesser, M.P. (2010). Nitrogen fixation and nitrogen transformations in marine symbioses. Trends in Microbiology 18, 455–463. 10.1016/j.tim.2010.07.001.

13. Coale, T.H., Loconte, V., Turk-Kubo, K.A., Vanslembrouck, B., Mak, W.K.E., Cheung, S., Ekman, A., Chen, J.-H., Hagino, K., Takano, Y., et al. (2024). Nitrogen-fixing organelle in a marine alga. Science 384, 217–222. 10.1126/science.adk1075.

14. Cornejo-Castillo, F.M., Inomura, K., Zehr, J.P., and Follows, M.J. (2024). Metabolic trade-offs constrain the cell size ratio in a nitrogen-fixing symbiosis. Cell 187, 1762–1768.e9. 10.1016/j.cell.2024.02.016.

15. Cabello, A.M., Turk-Kubo, K.A., Hayashi, K., Jacobs, L., Kudela, R.M., and Zehr, J.P. (2020). Unexpected presence of the nitrogen-fixing symbiotic cyanobacterium UCYN-A in Monterey Bay, California. Journal of Phycology 56, 1521–1533. 10.1111/jpy.13045.

16. Zehr, J.P., and Capone, D.G. (2020). Changing perspectives in marine nitrogen fixation. Science 368, eaay9514. 10.1126/science.aay9514.

17. Martínez-Pérez, C., Mohr, W., Löscher, C.R., Dekaezemacker, J., Littmann, S., Yilmaz, P., Lehnen, N., Fuchs, B.M., Lavik, G., Schmitz, R.A., et al. (2016). The small unicellular diazotrophic symbiont, UCYN-A, is a key player in the marine nitrogen cycle. Nat Microbiol 1, 16163. 10.1038/nmicrobiol.2016.163.

18. Tripp, H.J., Bench, S.R., Turk, K.A., Foster, R.A., Desany, B.A., Niazi, F., Affourtit, J.P., and Zehr, J.P. (2010). Metabolic streamlining in an open-ocean nitrogen-fixing cyanobacterium. Nature 464, 90–94. 10.1038/nature08786.

19. Bombar, D., Heller, P., Sanchez-Baracaldo, P., Carter, B.J., and Zehr, J.P. (2014). Comparative genomics reveals surprising divergence of two closely related strains of uncultivated UCYN-A cyanobacteria. The ISME Journal 8, 2530–2542. 10.1038/ismej.2014.167.

20. Cornejo-Castillo, F.M., Cabello, A.M., Salazar, G., Sánchez-Baracaldo, P., Lima-Mendez, G., Hingamp, P., Alberti, A., Sunagawa, S., Bork, P., De Vargas, C., et al. (2016). Cyanobacterial symbionts diverged in the late Cretaceous towards lineage-specific nitrogen fixation factories in single-celled phytoplankton. Nat Commun 7, 11071. 10.1038/ncomms11071.

21. Verma, S.K., and White, J.F. (2025). From ‘nitrosome’ to ‘nitroplast’: stages in the evolution of nitrogen-fixing organelles from free-living diazotrophs. Symbiosis 95, 29–34. 10.1007/s13199-025-01033-6.

22. Muñoz-Marín, M.D.C., Shilova, I.N., Shi, T., Farnelid, H., Cabello, A.M., and Zehr, J.P. (2019). The Transcriptional Cycle Is Suited to Daytime N_2_ Fixation in the Unicellular Cyanobacterium “ *Candidatus* Atelocyanobacterium thalassa” (UCYN-A). mBio 10, e02495–18. 10.1128/mBio.02495-18.

23. Church, M.J., Short, C.M., Jenkins, B.D., Karl, D.M., and Zehr, J.P. (2005). Temporal Patterns of Nitrogenase Gene (*nifH*) Expression in the Oligotrophic North Pacific Ocean. Appl Environ Microbiol 71, 5362–5370. 10.1128/AEM.71.9.5362-5370.2005.

24. Herman, E.M., and Sweeney, B.M. (1975). Circadian rhythm of chloroplast ultrastructure in Gonyaulax polyedra, concentric organization around a central cluster of ribosomes. Journal of Ultrastructure Research 50, 347–354. 10.1016/S0022-5320(75)80065-7.

25. Schmitt, K., Grimm, A., Dallmann, R., Oettinghaus, B., Restelli, L.M., Witzig, M., Ishihara, N., Mihara, K., Ripperger, J.A., Albrecht, U., et al. (2018). Circadian Control of DRP1 Activity Regulates Mitochondrial Dynamics and Bioenergetics. Cell Metabolism 27, 657–666.e5. 10.1016/j.cmet.2018.01.011.

26. Noordally, Z.B., Ishii, K., Atkins, K.A., Wetherill, S.J., Kusakina, J., Walton, E.J., Kato, M., Azuma, M., Tanaka, K., Hanaoka, M., et al. (2013). Circadian Control of Chloroplast Transcription by a Nuclear-Encoded Timing Signal. Science 339, 1316–1319. 10.1126/science.1230397.

27. Gradoville, M.R., Cabello, A.M., Wilson, S.T., Turk-Kubo, K.A., Karl, D.M., and Zehr, J.P. (2021). Light and depth dependency of nitrogen fixation by the non-photosynthetic, symbiotic cyanobacterium UCYN-A. Environmental Microbiology 23, 4518–4531. 10.1111/1462-2920.15645.

28. Landa, M., Turk-Kubo, K.A., Cornejo-Castillo, F.M., Henke, B.A., and Zehr, J.P. (2021). Critical Role of Light in the Growth and Activity of the Marine N2-Fixing UCYN-A Symbiosis. Front. Microbiol. 12, 666739. 10.3389/fmicb.2021.666739.

29. Moulin, S.L.Y., Frail, S., Braukmann, T., Doenier, J., Steele-Ogus, M., Marks, J.C., Mills, M.M., and Yeh, E. (2024). The endosymbiont of *Epithemia clementina* is specialized for nitrogen fixation within a photosynthetic eukaryote. ISME Communications 4, ycae055. 10.1093/ismeco/ycae055.

30. Hagino, K., Onuma, R., Kawachi, M., and Horiguchi, T. (2013). Discovery of an Endosymbiotic Nitrogen-Fixing Cyanobacterium UCYN-A in Braarudosphaera bigelowii (Prymnesiophyceae). PLoS ONE 8, e81749. 10.1371/journal.pone.0081749.

31. Suzuki, S., Kawachi, M., Tsukakoshi, C., Nakamura, A., Hagino, K., Inouye, I., and Ishida, K. (2021). Unstable Relationship Between Braarudosphaera bigelowii (= Chrysochromulina parkeae) and Its Nitrogen-Fixing Endosymbiont. Front. Plant Sci. 12, 749895. 10.3389/fpls.2021.749895.

32. Mak, E.W.K., Turk-Kubo, K.A., Caron, D.A., Harbeitner, R.C., Magasin, J.D., Coale, T.H., Hagino, K., Takano, Y., Nishimura, T., Adachi, M., et al. (2024). Phagotrophy in the nitrogen-fixing haptophyte *Braarudosphaera bigelowii* . Environ Microbiol Rep 16, e13312. 10.1111/1758-2229.13312.

33. Cabello, A.M., Cornejo-Castillo, F.M., Raho, N., Blasco, D., Vidal, M., Audic, S., De Vargas, C., Latasa, M., Acinas, S.G., and Massana, R. (2016). Global distribution and vertical patterns of a prymnesiophyte–cyanobacteria obligate symbiosis. The ISME Journal 10, 693–706. 10.1038/ismej.2015.147.

34. Kaplan, M., Nicolas, W.J., Zhao, W., Carter, S.D., Metskas, L.A., Chreifi, G., Ghosal, D., and Jensen, G.J. (2021). In Situ Imaging and Structure Determination of Biomolecular Complexes Using Electron Cryo-Tomography. In cryoEM Methods in Molecular Biology., T. Gonen and B. L. Nannenga, eds. (Springer US), pp. 83–111. 10.1007/978-1-0716-0966-8_4.

35. Marshall, A.G., Damo, S.M., and Hinton, A. (2023). Revisiting focused ion beam scanning electron microscopy. Trends in Biochemical Sciences 48, 585–586. 10.1016/j.tibs.2023.02.005.

36. Bousquet, L., Fainsod, S., Decelle, J., Murik, O., Chevalier, F., Gallet, B., Templin, R., Schwab, Y., Avrahami, Y., Koplovitz, G., et al. (2025). Life cycle and morphogenetic differentiation in heteromorphic cell types of a cosmopolitan marine microalga. New Phytologist 245, 1969–1984. 10.1111/nph.20360.

37. Wheeler, G.L., Sturm, D., and Langer, G. (2023). *Gephyrocapsa huxleyi* (*Emiliania huxleyi*) as a model system for coccolithophore biology. Journal of Phycology 59, 1123–1129. 10.1111/jpy.13404.

38. Uwizeye, C., Mars Brisbin, M., Gallet, B., Chevalier, F., LeKieffre, C., Schieber, N.L., Falconet, D., Wangpraseurt, D., Schertel, L., Stryhanyuk, H., et al. (2021). Cytoklepty in the plankton: A host strategy to optimize the bioenergetic machinery of endosymbiotic algae. Proc. Natl. Acad. Sci. U.S.A. 118, e2025252118. 10.1073/pnas.2025252118.

39. Godrijan, J., Drapeau, D.T., and Balch, W.M. (2022). Osmotrophy of dissolved organic carbon by coccolithophores in darkness. New Phytologist 233, 781–794. 10.1111/nph.17819.

40. Mocaer, K., Mizzon, G., Gunkel, M., Halavatyi, A., Steyer, A., Oorschot, V., Schorb, M., Le Kieffre, C., Yee, D.P., Chevalier, F., et al. (2023). Targeted volume correlative light and electron microscopy of an environmental marine microorganism. Journal of Cell Science 136, jcs261355. 10.1242/jcs.261355.

41. Zehr, J.P., Bench, S.R., Carter, B.J., Hewson, I., Niazi, F., Shi, T., Tripp, H.J., and Affourtit, J.P. (2008). Globally Distributed Uncultivated Oceanic N_2_ -Fixing Cyanobacteria Lack Oxygenic Photosystem II. Science 322, 1110–1112. 10.1126/science.1165340.

42. Thompson, A.W., Foster, R.A., Krupke, A., Carter, B.J., Musat, N., Vaulot, D., Kuypers, M.M.M., and Zehr, J.P. (2012). Unicellular Cyanobacterium Symbiotic with a Single-Celled Eukaryotic Alga. Science 337, 1546–1550. 10.1126/science.1222700.

43. Austin, J.R., Frost, E., Vidi, P.-A., Kessler, F., and Staehelin, L.A. (2006). Plastoglobules Are Lipoprotein Subcompartments of the Chloroplast That Are Permanently Coupled to Thylakoid Membranes and Contain Biosynthetic Enzymes. Plant Cell 18, 1693–1703. 10.1105/tpc.105.039859.

44. Blum, T.B., Hahn, A., Meier, T., Davies, K.M., and Kühlbrandt, W. (2019). Dimers of mitochondrial ATP synthase induce membrane curvature and self-assemble into rows. Proc. Natl. Acad. Sci. U.S.A. 116, 4250–4255. 10.1073/pnas.1816556116.

45. Voeltz, G.K., Sawyer, E.M., Hajnóczky, G., and Prinz, W.A. (2024). Making the connection: How membrane contact sites have changed our view of organelle biology. Cell 187, 257–270. 10.1016/j.cell.2023.11.040.

46. Nowack, E.C.M., Melkonian, M., and Glöckner, G. (2008). Chromatophore Genome Sequence of Paulinella Sheds Light on Acquisition of Photosynthesis by Eukaryotes. Current Biology 18, 410–418. 10.1016/j.cub.2008.02.051.

47. Delaye, L., Valadez-Cano, C., and Pérez-Zamorano, B. (2016). How Really Ancient Is Paulinella Chromatophora? PLoS Curr. 10.1371/currents.tol.e68a099364bb1a1e129a17b4e06b0c6b.

48. Macorano, L., and Nowack, E.C.M. (2021). Paulinella chromatophora. Current Biology 31, R1024–R1026. 10.1016/j.cub.2021.07.028.

49. Howard, J.B., and Rees, D.C. (1996). Structural Basis of Biological Nitrogen Fixation. Chem. Rev. 96, 2965–2982. 10.1021/cr9500545.

50. Falkowski, P.G., and Stone, D.P. (1975). Nitrate uptake in marine phytoplankton: Energy sources and the interaction with carbon fixation. Mar. Biol. 32, 77–84. 10.1007/BF00395161.

51. Glibert, P.M., Wilkerson, F.P., Dugdale, R.C., Raven, J.A., Dupont, C.L., Leavitt, P.R., Parker, A.E., Burkholder, J.M., and Kana, T.M. (2016). Pluses and minuses of ammonium and nitrate uptake and assimilation by phytoplankton and implications for productivity and community composition, with emphasis on nitrogen-enriched conditions: Pluses and minuses of NH4+ *and* NO3−. Limnol. Oceanogr. 61, 165–197. 10.1002/lno.10203.

52. Mills, M.M., Turk-Kubo, K.A., Van Dijken, G.L., Henke, B.A., Harding, K., Wilson, S.T., Arrigo, K.R., and Zehr, J.P. (2020). Unusual marine cyanobacteria/haptophyte symbiosis relies on N2 fixation even in N-rich environments. The ISME Journal 14, 2395–2406. 10.1038/s41396-020-0691-6.

53. Csordás, G., Renken, C., Várnai, P., Walter, L., Weaver, D., Buttle, K.F., Balla, T., Mannella, C.A., and Hajnóczky, G. (2006). Structural and functional features and significance of the physical linkage between ER and mitochondria. The Journal of Cell Biology 174, 915–921. 10.1083/jcb.200604016.

54. Gould, S.B., Waller, R.F., and McFadden, G.I. (2008). Plastid Evolution. Annu. Rev. Plant Biol. 59, 491–517. 10.1146/annurev.arplant.59.032607.092915.

55. MacLeod, A.I., Knopp, M.R., and Gould, S.B. (2024). A mysterious cloak: the peptidoglycan layer of algal and plant plastids. Protoplasma 261, 173–178. 10.1007/s00709-023-01886-y.

56. Samuel, A.D., Petersen, J.D., and Reese, T.S. (2001). Envelope structure of Synechococcus sp. WH8113, a nonflagellated swimming cyanobacterium. BMC Microbiol 1, 4. 10.1186/1471-2180-1-4.

57. Ting, C.S., Hsieh, C., Sundararaman, S., Mannella, C., and Marko, M. (2007). Cryo-Electron Tomography Reveals the Comparative Three-Dimensional Architecture of *Prochlorococcus*, a Globally Important Marine Cyanobacterium. J Bacteriol 189, 4485–4493. 10.1128/JB.01948-06.

58. Al-Amoudi, A., Chang, J.-J., Leforestier, A., McDowall, A., Salamin, L.M., Norlén, L.P.O., Richter, K., Blanc, N.S., Studer, D., and Dubochet, J. (2004). Cryo-electron microscopy of vitreous sections. EMBO J 23, 3583–3588. 10.1038/sj.emboj.7600366.

59. Bergman, B., Sandh, G., Lin, S., Larsson, J., and Carpenter, E.J. (2013). *Trichodesmium* – a widespread marine cyanobacterium with unusual nitrogen fixation properties. FEMS Microbiol Rev 37, 286–302. 10.1111/j.1574-6976.2012.00352.x.

60. Bale, N.J., Villareal, T.A., Hopmans, E.C., Brussaard, C.P.D., Besseling, M., Dorhout, D., Sinninghe Damsté, J.S., and Schouten, S. (2018). C_5_ glycolipids of heterocystous cyanobacteria track symbiont abundance in the diatom *Hemiaulus hauckii* across the tropical North Atlantic. Biogeosciences 15, 1229–1241. 10.5194/bg-15-1229-2018.

61. Schouten, S., Villareal, T.A., Hopmans, E.C., Mets, A., Swanson, K.M., and Sinninghe Damsté, J.S. (2013). Endosymbiotic heterocystous cyanobacteria synthesize different heterocyst glycolipids than free-living heterocystous cyanobacteria. Phytochemistry 85, 115–121. 10.1016/j.phytochem.2012.09.002.

62. Takano, H., and Takechi, K. (2010). Plastid peptidoglycan. Biochimica et Biophysica Acta (BBA) - General Subjects 1800, 144–151. 10.1016/j.bbagen.2009.07.020.

63. Limpens, E. (2019). Extracellular membranes in symbiosis. Nature Plants 5, 131–132. 10.1038/s41477-019-0370-7.

64. Ivanov, S., Austin, J., Berg, R.H., and Harrison, M.J. (2019). Extensive membrane systems at the host–arbuscular mycorrhizal fungus interface. Nature Plants 5, 194–203. 10.1038/s41477-019-0364-5.

65. Roth, R., Hillmer, S., Funaya, C., Chiapello, M., Schumacher, K., Lo Presti, L., Kahmann, R., and Paszkowski, U. (2019). Arbuscular cell invasion coincides with extracellular vesicles and membrane tubules. Nature Plants 5, 204–211. 10.1038/s41477-019-0365-4.

66. Biller, S.J., Schubotz, F., Roggensack, S.E., Thompson, A.W., Summons, R.E., and Chisholm, S.W. (2014). Bacterial Vesicles in Marine Ecosystems. Science 343, 183–186. 10.1126/science.1243457.

67. Biller, S.J., Lundeen, R.A., Hmelo, L.R., Becker, K.W., Arellano, A.A., Dooley, K., Heal, K.R., Carlson, L.T., Van Mooy, B.A.S., Ingalls, A.E., et al. (2022). *Prochlorococcus* extracellular vesicles: molecular composition and adsorption to diverse microbes. Environmental Microbiology 24, 420–435. 10.1111/1462-2920.15834.

68. Pardo, Y.A., Florez, C., Baker, K.M., Schertzer, J.W., and Mahler, G.J. (2015). Detection of outer membrane vesicles in *Synechocystis* PCC 6803. FEMS Microbiology Letters 362, fnv163. 10.1093/femsle/fnv163.

69. Flores, E., Romanovicz, D.K., Nieves-Morión, M., Foster, R.A., and Villareal, T.A. (2022). Adaptation to an Intracellular Lifestyle by a Nitrogen-Fixing, Heterocyst-Forming Cyanobacterial Endosymbiont of a Diatom. Front. Microbiol. 13, 799362. 10.3389/fmicb.2022.799362.

70. McQuaid, J.B., Kustka, A.B., Oborník, M., Horák, A., McCrow, J.P., Karas, B.J., Zheng, H., Kindeberg, T., Andersson, A.J., Barbeau, K.A., et al. (2018). Carbonate-sensitive phytotransferrin controls high-affinity iron uptake in diatoms. Nature 555, 534–537. 10.1038/nature25982.

71. Turnšek, J., Brunson, J.K., Viedma, M.D.P.M., Deerinck, T.J., Horák, A., Oborník, M., Bielinski, V.A., and Allen, A.E. (2021). Proximity proteomics in a marine diatom reveals a putative cell surface-to-chloroplast iron trafficking pathway. eLife 10, e52770. 10.7554/eLife.52770.

72. Kaplan, M., Oikonomou, C.M., Wood, C.R., Chreifi, G., Ghosal, D., Dobro, M.J., Yao, Q., Pal, R.R., Baidya, A.K., Liu, Y., et al. (2022). Discovery of a Novel Inner Membrane-Associated Bacterial Structure Related to the Flagellar Type III Secretion System. J Bacteriol, e00144–22. 10.1128/jb.00144-22.

73. Pujhari, S., Heebner, J., Raumann, E., Zhong, T., Rasgon, J.L., Swulius, M.T., Shaffer, C.L., and Kaplan, M. (2025). *In situ* architecture of the endosymbiont *Wolbachia pipientis*. Preprint at Microbiology, 10.1101/2025.08.29.673095.

74. Wietrzynski, W., Lamm, L., Wood, W.H., Loukeri, M.-J., Malone, L., Peng, T., Johnson, M.P., and Engel, B.D. (2025). Molecular architecture of thylakoid membranes within intact spinach chloroplasts. Preprint, 10.7554/eLife.105496.2.

75. Rose, K., Herrmann, E., Kakudji, E., Lizarrondo, J., Celebi, A.Y., Wilfling, F., Lewis, S.C., and Hurley, J.H. (2025). In situ cryo-ET visualization of mitochondrial depolarization and mitophagic engulfment. Proc. Natl. Acad. Sci. U.S.A. 122, e2511890122. 10.1073/pnas.2511890122.

76. Gray, M.W. (2012). Mitochondrial Evolution. Cold Spring Harbor Perspectives in Biology 4, a011403–a011403. 10.1101/cshperspect.a011403.

77. Shi, L.-X., and Theg, S.M. (2013). The chloroplast protein import system: From algae to trees. Biochimica et Biophysica Acta (BBA) - Molecular Cell Research 1833, 314–331. 10.1016/j.bbamcr.2012.10.002.

78. Chacinska, A., Koehler, C.M., Milenkovic, D., Lithgow, T., and Pfanner, N. (2009). Importing Mitochondrial Proteins: Machineries and Mechanisms. Cell 138, 628–644. 10.1016/j.cell.2009.08.005.

79. Good, A. (2018). Toward nitrogen-fixing plants. Science 359, 869–870. 10.1126/science.aas8737.

80. Guillard, R.R.L., and Ryther, J.H. (1962). STUDIES OF MARINE PLANKTONIC DIATOMS: I. CYCLOTELLA NANA HUSTEDT, AND DETONULA CONFERVACEA (CLEVE) GRAN. Can. J. Microbiol. 8, 229–239. 10.1139/m62-029.

81. Rao, A.K., Yee, D., Chevalier, F., LeKieffre, C., Pavie, M., Olivetta, M., Dudin, O., Gallet, B., Hehenberger, E., Seifi, M., et al. (2025). Hijacking and integration of algal plastids and mitochondria in a polar planktonic host. Current Biology 35, 2509–2523.e7. 10.1016/j.cub.2025.03.076.

82. Gallet, B., Moriscot, C., Schoehn, G., and Decelle, J. (2024). Cryo-fixation and resin embedding of biological samples for electron microscopy and chemical imaging. Preprint, 10.17504/protocols.io.bp2l62kndgqe/v1.

83. Ollion, J., Cochennec, J., Loll, F., Escudé, C., and Boudier, T. (2013). TANGO: a generic tool for high-throughput 3D image analysis for studying nuclear organization. Bioinformatics 29, 1840–1841. 10.1093/bioinformatics/btt276.

84. Fedorov, A., Beichel, R., Kalpathy-Cramer, J., Finet, J., Fillion-Robin, J.-C., Pujol, S., Bauer, C., Jennings, D., Fennessy, F., Sonka, M., et al. (2012). 3D Slicer as an image computing platform for the Quantitative Imaging Network. Magnetic Resonance Imaging 30, 1323–1341. 10.1016/j.mri.2012.05.001.

85. Ayachit, U. (2015). The ParaView Guide: A Parallel Visualization Application (Kitware, Inc., 28 Corporate Drive, Clifton Park, NY, United States).

86. Leisch, N., Baars, S., Beavis, T., Bertucci, P., Bhickta, C., Bonadonna, M., Brannon, C., Burgués-Palau, L., Cherek, P., Chevalier, F., et al. (2026). An Advanced Mobile Laboratory to enable field-based microbial ecology and cell biology across scales. Preprint at Cell Biology, 10.64898/2026.02.23.707475.

87. Ronchi, P., Mizzon, G., Machado, P., D’Imprima, E., Best, B.T., Cassella, L., Schnorrenberg, S., Montero, M.G., Jechlinger, M., Ephrussi, A., et al. (2021). High-precision targeting workflow for volume electron microscopy. Journal of Cell Biology 220, e202104069. 10.1083/jcb.202104069.

88. Hennies, J., Lleti, J.M.S., Schieber, N.L., Templin, R.M., Steyer, A.M., and Schwab, Y. (2020). AMST: Alignment to Median Smoothed Template for Focused Ion Beam Scanning Electron Microscopy Image Stacks. Sci Rep 10, 2004. 10.1038/s41598-020-58736-7.

89. Hagen, W.J.H., Wan, W., and Briggs, J.A.G. (2017). Implementation of a cryo-electron tomography tilt-scheme optimized for high resolution subtomogram averaging. J. Struct. Biol. 197, 191–198. 10.1016/j.jsb.2016.06.007.

90. Kremer, J.R., Mastronarde, D.N., and McIntosh, J.R. (1996). Computer visualization of three-dimensional image data using IMOD. J. Struct. Biol. 116, 71–76. 10.1006/jsbi.1996.0013.

91. Nicastro, D. (2006). The Molecular Architecture of Axonemes Revealed by Cryoelectron Tomography. Science 313, 944–948. 10.1126/science.1128618.

92. Castaño-Díez, D., Kudryashev, M., Arheit, M., and Stahlberg, H. (2012). Dynamo: a flexible, user-friendly development tool for subtomogram averaging of cryo-EM data in high-performance computing environments. J. Struct. Biol. 178, 139–151. 10.1016/j.jsb.2011.12.017.

93. Scaramuzza, S., and Castaño-Díez, D. (2021). Step-by-step guide to efficient subtomogram averaging of virus-like particles with Dynamo. PLoS Biol 19, e3001318. 10.1371/journal.pbio.3001318.

94. Coale, T., Loconte, V., Turk-Kubo, K., Vanslembrouck, B., Mak, W.K.E., Cheung, S., Ekman, A., Chen, J.-H., Hagino, K., Takano, Y., et al. (2024). Data from: Nitrogen fixing organelle in a marine alga. Version 16 (Dryad). 10.5061/DRYAD.2Z34TMPTF.

95. Ritchie, M.E., Phipson, B., Wu, D., Hu, Y., Law, C.W., Shi, W., and Smyth, G.K. (2015). limma powers differential expression analyses for RNA-sequencing and microarray studies. Nucleic Acids Research 43, e47–e47. 10.1093/nar/gkv007.

