## Supplemental Figures for "Diel remodeling and cellular integration of the nitroplast"

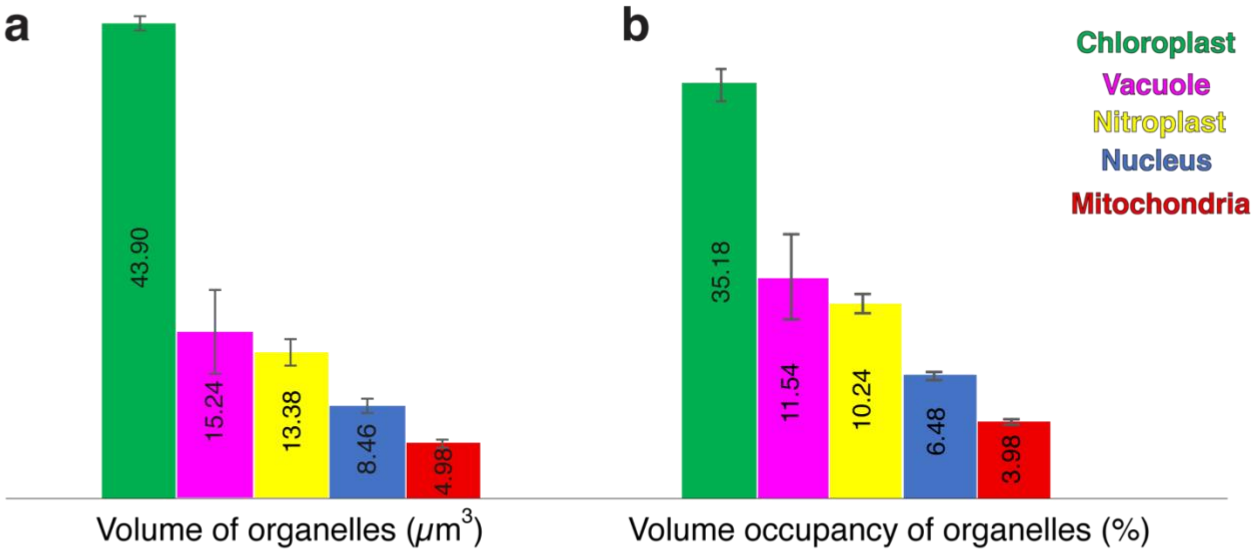


**Figure S1 | Organelles morphometric measurements.**

Organelle morphometrics derived from FIB–SEM reconstructions: **a,** absolute volumes ($\mu m^3$) and **b,** fractional volume occupancy (% of total cell volume).

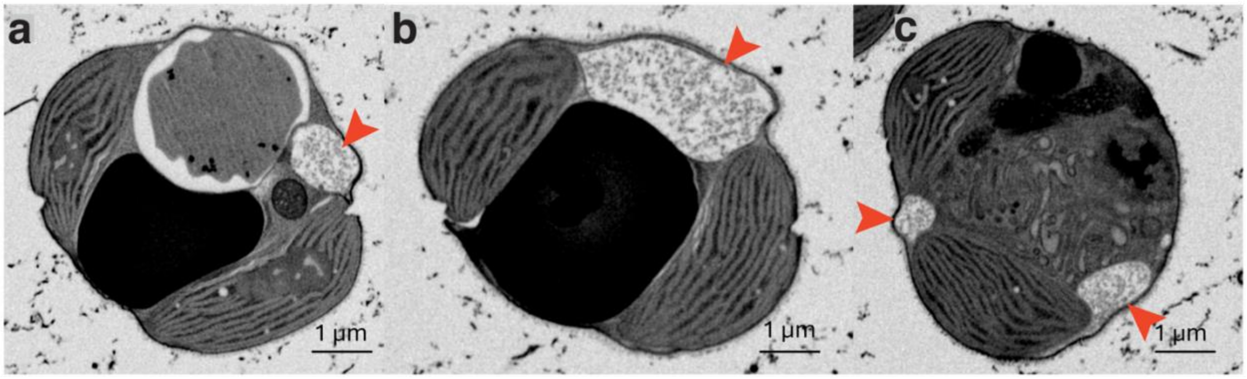


**Figure S2 | Tubular meshwork adjacent to the nitroplast in cultured cells.**

Slices through a FIB–SEM stack of a cultured, motile *B. bigelowii* cell at multiple z-heights highlighting a membrane-bound compartment containing a mesh of tubules (red arrows).

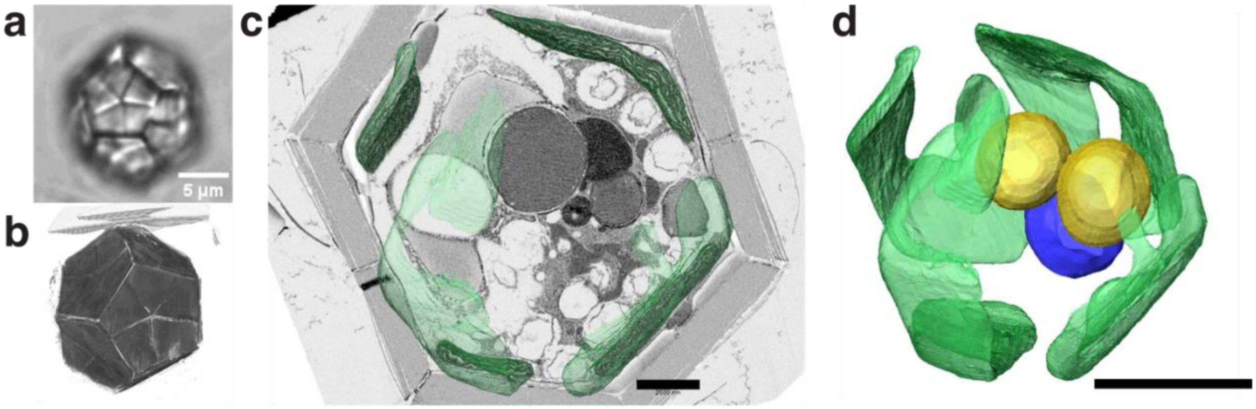


**Figure S3 | FIB-SEM imaging of the non-motile, calcified stage of *B. bigelowii* collected in the natural environment.**

**a,** Light microscopy of a resin-embedded calcified cell (~15 µm in diameter) with thick pentaliths. **b,** Volumetric rendering of a second cell from the FIB–SEM dataset. **c-d**, FIB-SEM imaging and 3D reconstruction of a third cell, highlighting four chloroplasts (green), two nitroplasts (yellow), and one nucleus (blue). Scale bar 2 µm (c), 5 µm (d).

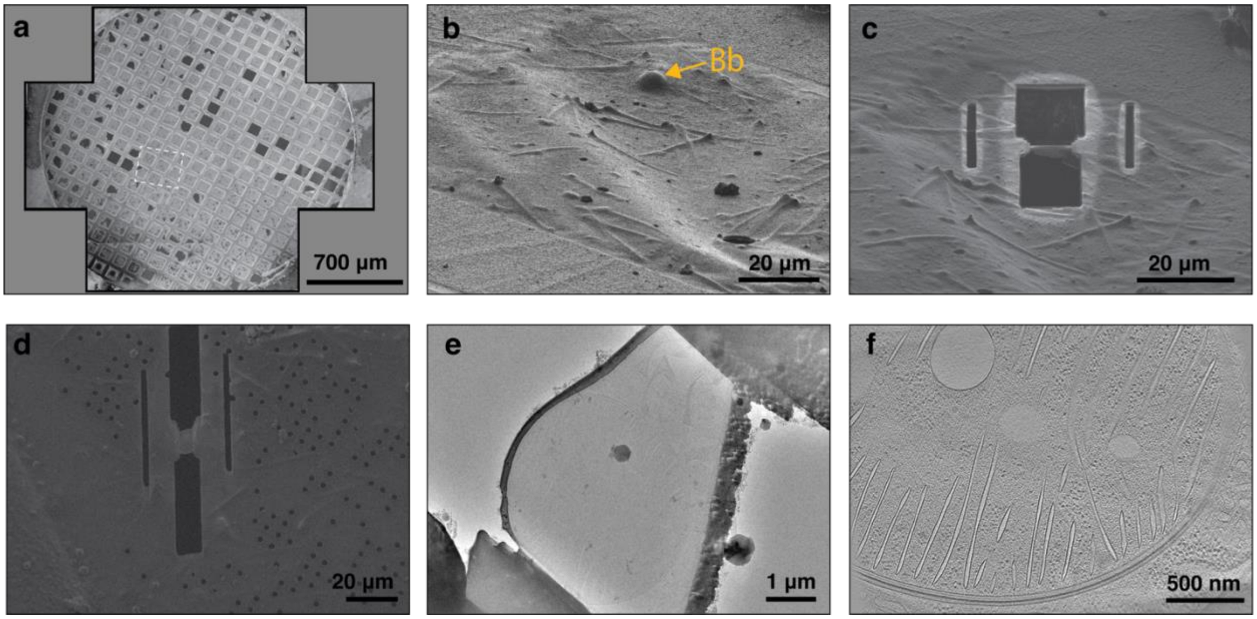


**Figure S4 | Cryo-FIB milling workflow for in situ cryo-electron tomography of *B. bigelowii*.**

**a,** Scanning electron microscopy (SEM) image of a grid containing plunge-frozen, cultured motile *B. bigelowii* cells. **b,** Targeted *B. bigelowii* cell (orange arrow) imaged prior to milling using an Aquilos 2 cryo-FIB microscope. **c,** Cryo-FIB milling procedure, showing gradual removal of cellular material above and below the lamella. **d,** Representative cryo-FIB–milled lamella. **e,** Two-dimensional cryo-TEM projection image of the lamella shown in d. **f,** Slice through a cryo-electron tomogram of the lamella shown in e, revealing a nitroplast.

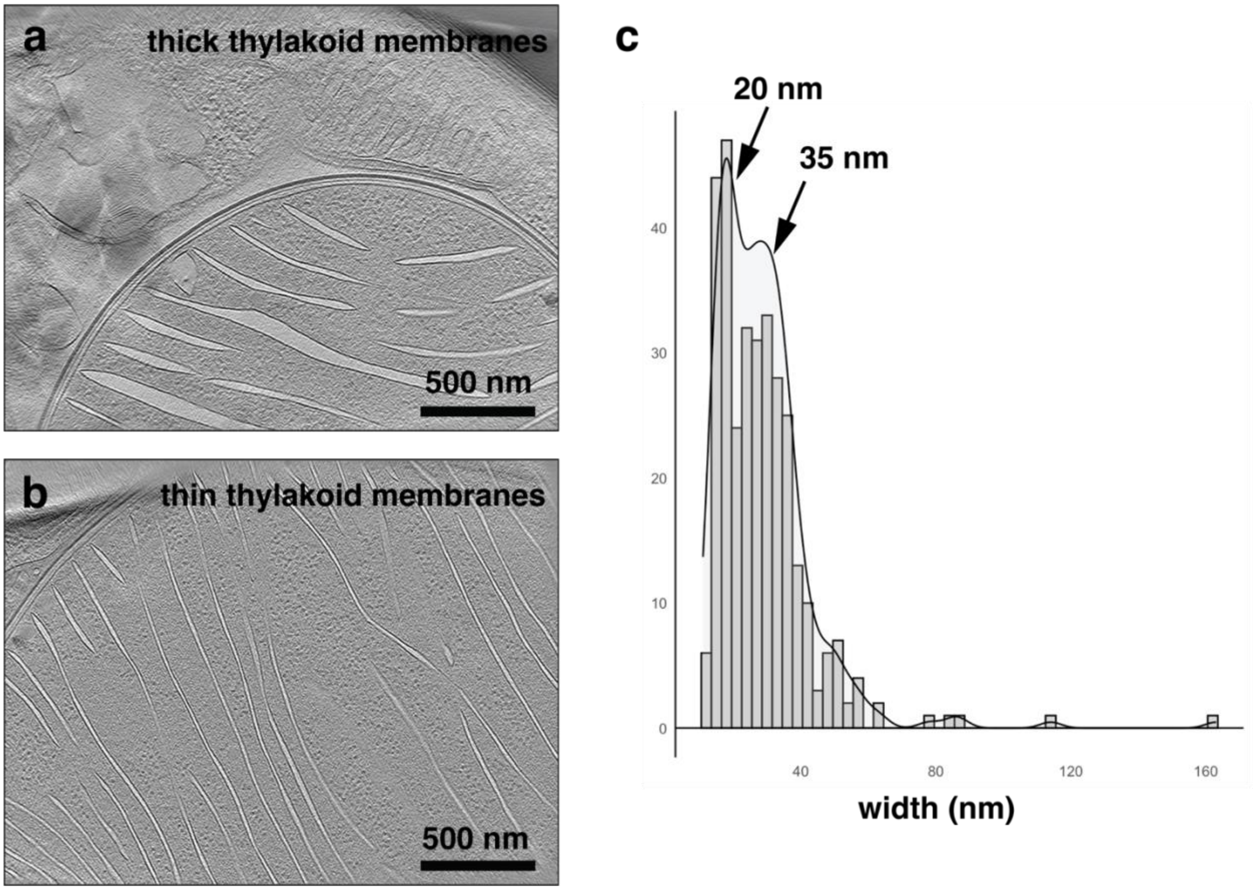


**Figure S5 | Thylakoid membrane thickness in morning nitroplasts.**

**a,b,** Cryo-electron tomogram slices of morning nitroplasts illustrating thylakoid membranes of variable thickness. **c,** Distribution of measured thylakoid thickness values in morning nitroplasts.

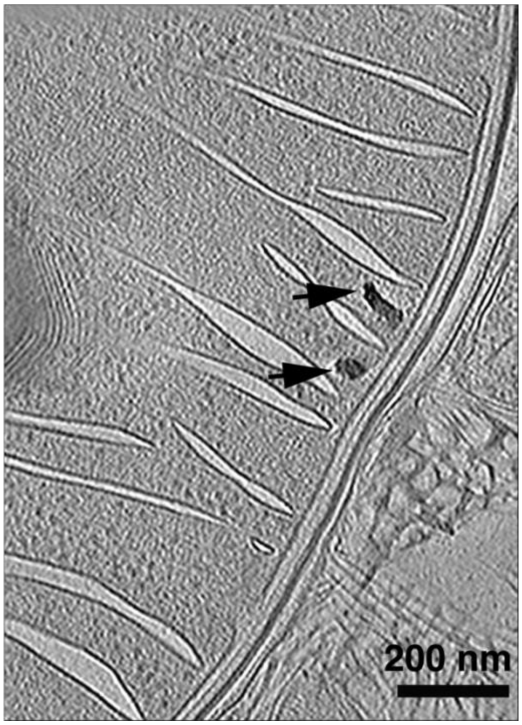


**Figure S6 | Electron-dense bodies near the nitroplast inner membrane.**
Slice through a cryo-electron tomogram of a morning nitroplast highlighting amorphous electron-dense structures located adjacent to the nitroplast inner membrane (IM) (black arrows).

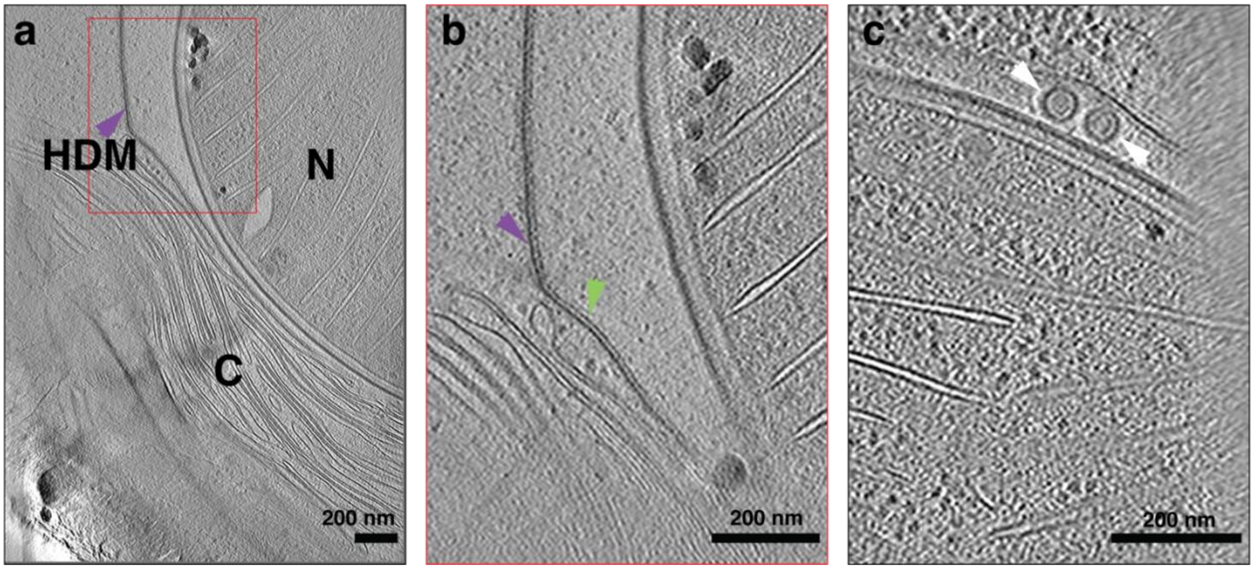


**Figure S7 | Host-derived layers and vesicles at the morning nitroplast interface.**

**a,** Cryo-electron tomogram slice of a morning nitroplast (N) adjacent to a chloroplast (C), highlighting the host-derived membrane (HDM; purple arrow) and the associated host granular layer (HGL; green arrow) positioned beneath it. Local regions where both layers drift from the nitroplast surface suggest physical coupling between HDM and HGL. **b,** Enlarged view of the boxed region in a. **c,** Cryo-electron tomogram slice showing spiky vesicles (white arrows) located between the nitroplast outer membrane (OM) and the HDM.

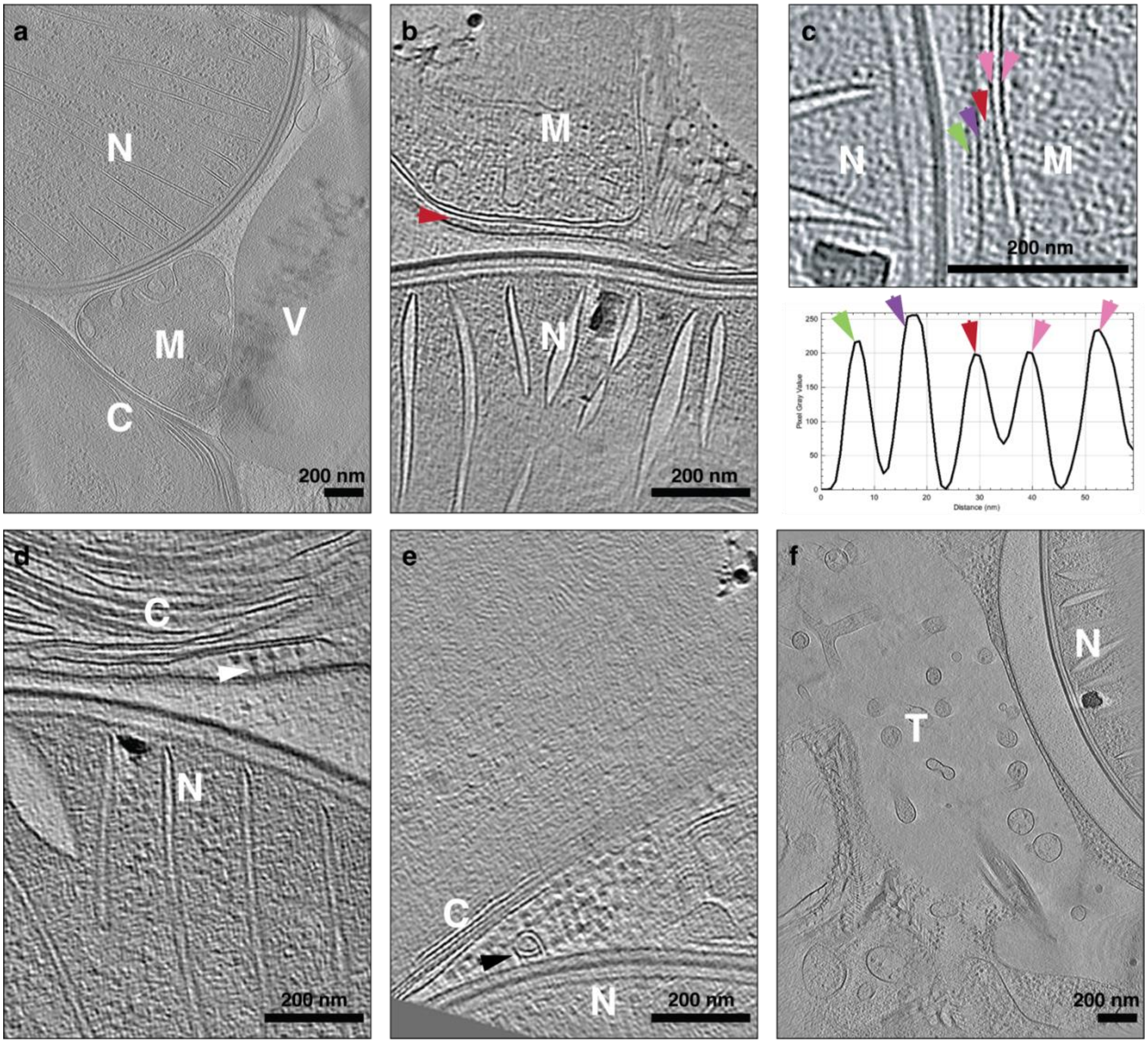


**Figure S8 | Membrane contact sites between the morning nitroplast and host organelles.**

**a,** Slice through a cryo-electron tomogram of a *B. bigelowii* cell showing the proximity of a morning nitroplast (N) to multiple host organelles, including mitochondrion (M), chloroplast (C), and vacuole (V). **b,** Slice through a cryo-electron tomogram highlighting an additional electron-dense layer (red arrow) positioned between the nitroplast host-derived membrane (HDM) and the mitochondrion (M). **c,** Top, slice through a cryo-electron tomogram showing the nitroplast–mitochondrion interface. Bottom, corresponding density profile across the contact site. Pink arrows indicate the mitochondrial inner and outer membranes; red arrow marks the additional interfacial layer; purple arrow indicates the HDM; green arrow indicates the host granular layer (HGL). **d,** Slice through a cryo-electron tomogram showing putative ribosomes (white arrows) associated with the HDM at the chloroplast (C)–nitroplast (N) interface. **e,** Slice through a cryo-electron tomogram showing a spiral structure (black arrow) in contact with the HDM at the chloroplast–nitroplast interface. **f,** Slice through a cryo-electron tomogram illustrating the interface between the nitroplast (N) and a membrane-bound tubular network (T).

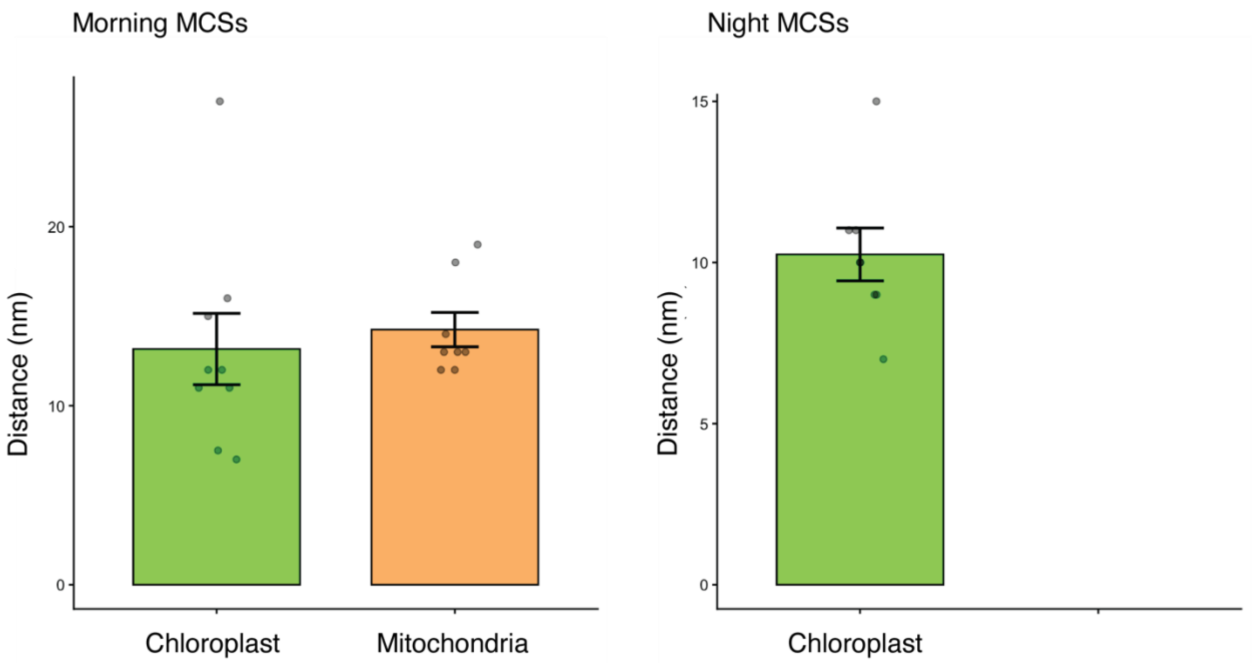


**Figure S9 | Spatial proximity of the nitroplast to other organelles.**
Distance between the nitroplast and chloroplasts (green) or mitochondria (orange) in morning (left) and night (right) cryo-tomograms. No mitochondria were observed in proximity to night nitroplasts in the analyzed datasets.

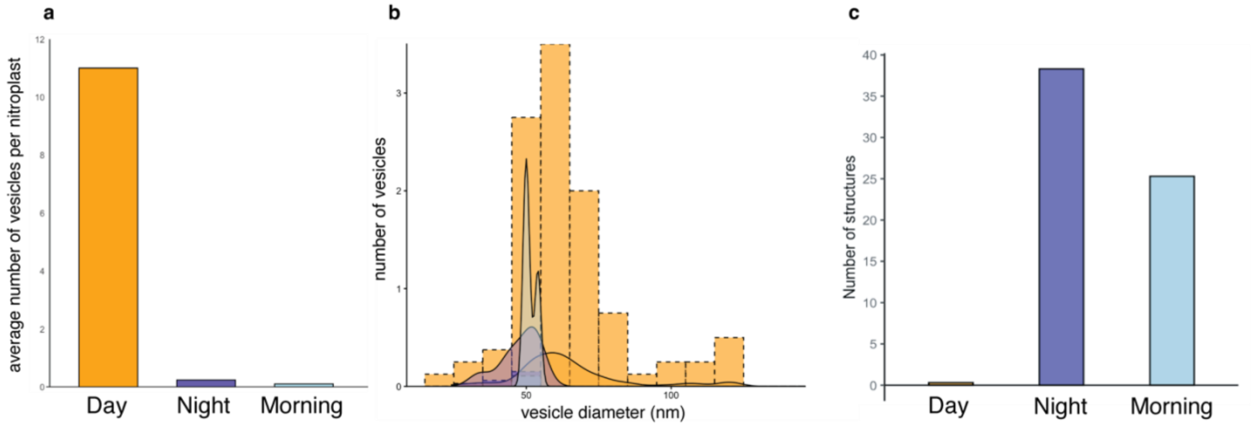


**Figure S10 | Quantification of vesicles and thylakoid-associated structures across the diel cycle.**

**a,** Average number of spiky vesicles per nitroplast in morning, day, and night samples. **b,** Diameter distribution of spiky vesicles across morning, day, and night datasets. **c,** Number of triangular (hat-like) thylakoid-associated complexes per nitroplast in morning, day, and night samples.

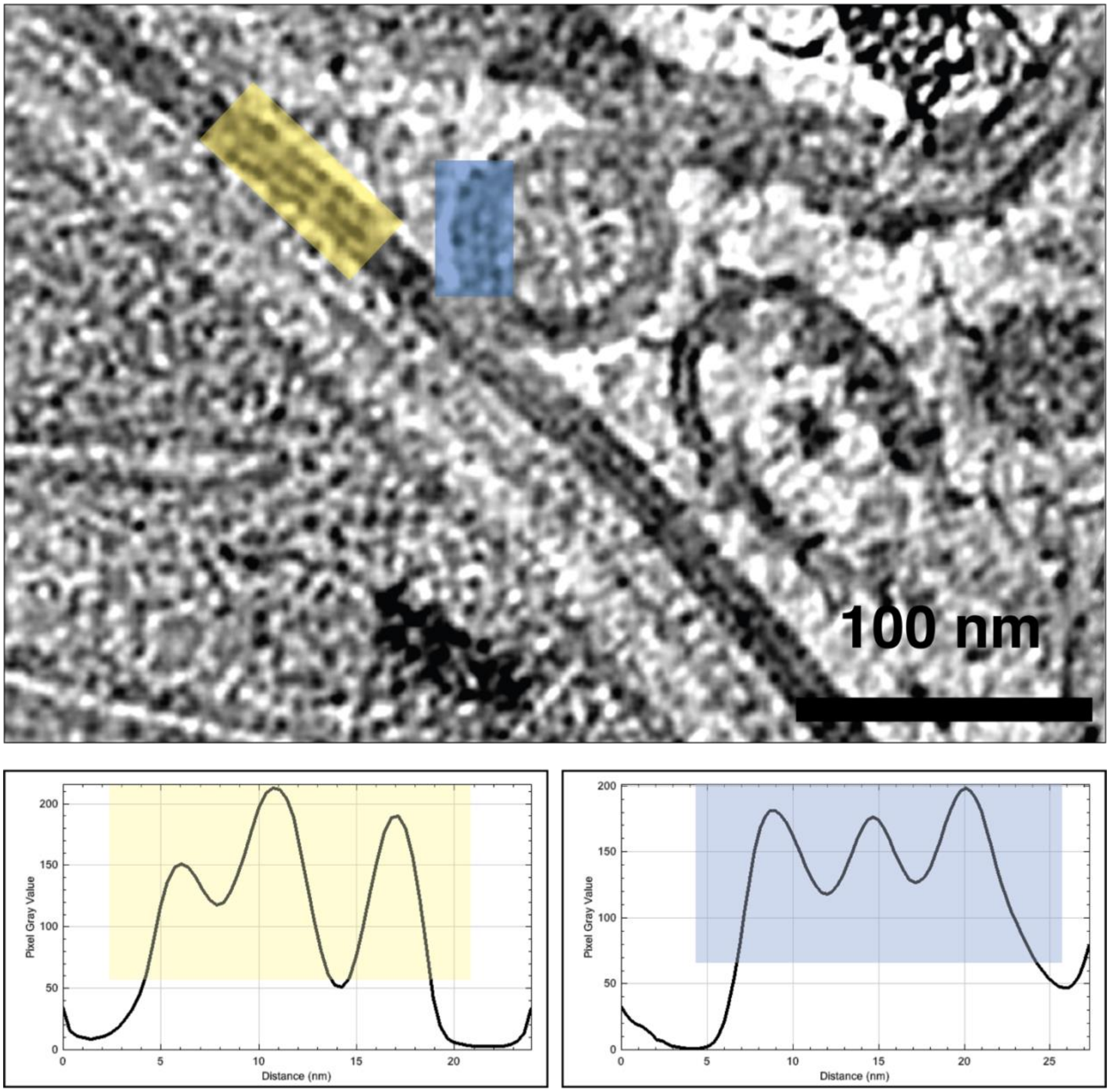


**Figure S11 | Membrane density profiles of spiky vesicles and the nitroplast outer membrane.**

Top: Cryo-electron tomogram slice of a day nitroplast (N) with a spiky vesicle attached to the nitroplast outer membrane (OM). Bottom left: Density profile of the nitroplast OM (yellow-boxed region in upper panel). Bottom right: Density profile of the spiky vesicle membrane (blue-boxed region). Both profiles display a similar three-peak pattern.

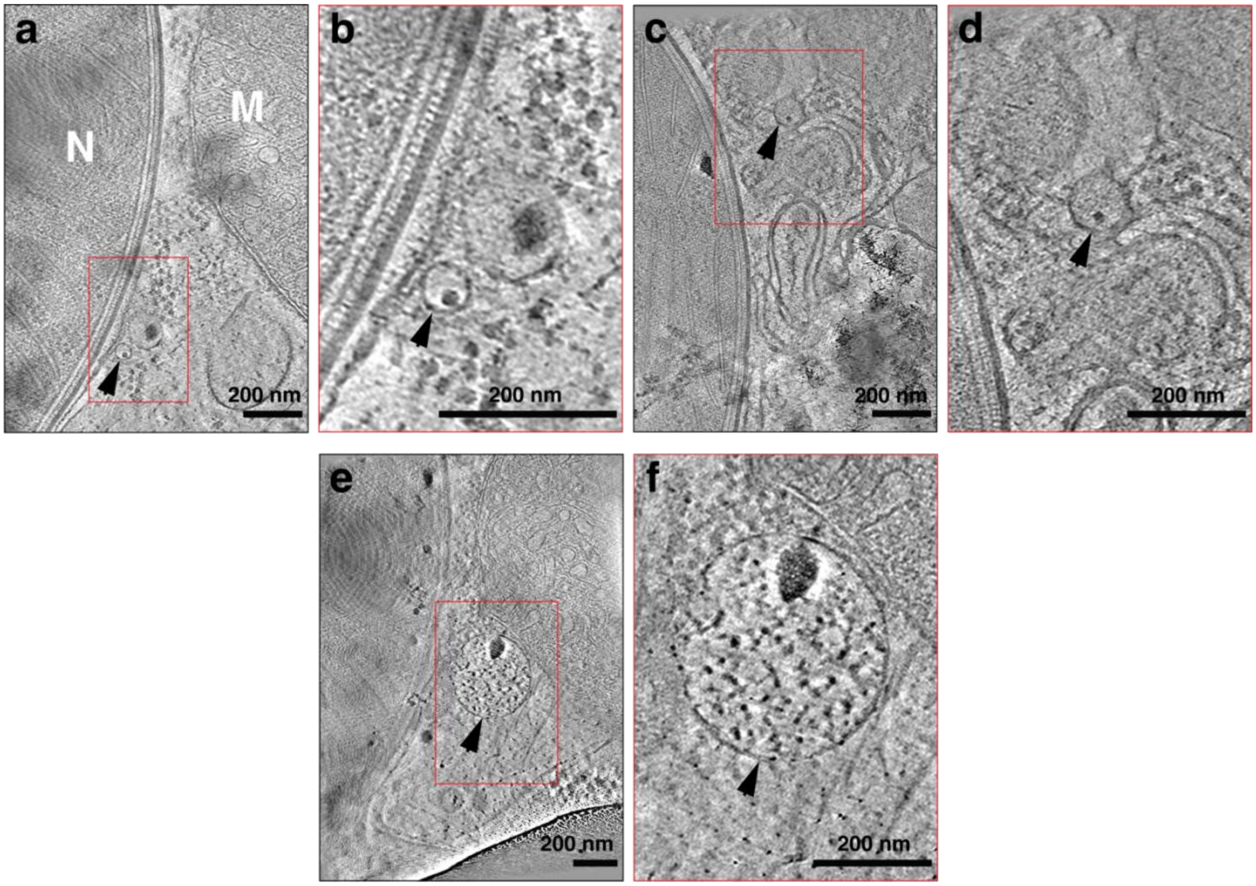


**Figure S12 | Smooth vesicles at the nitroplast–host interface during the day.**

**a–f,** Cryo-electron tomogram slices of day nitroplasts showing smooth vesicles containing electron-dense cargo (black arrows) at the nitroplast–host interface. Panels b, d, f are enlarged views of the boxed regions in a, c, e, respectively.
